# eDNA sampling systems for salmon ecosystem monitoring

**DOI:** 10.1101/2024.09.11.610878

**Authors:** Christoph M. Deeg, Robert G. Saunders, Christopher Tam, Karia Kaukinen, Shaorong Li, Arthur L Bass, Uu-a-thluk Fisheries, Kristina M. Miller

## Abstract

Environmental DNA (eDNA) is transforming the way aquatic ecosystems are monitored and managed by scientists, resource managers, ENGOs, First Nations communities, and citizen scientists alike. However, the lack of sampling systems enabling high filtration volumes and rapid sample collection in the field have thus far hindered broad scale eDNA studies in the ocean specifically for small and medium scale organizations. To overcome these challenges, several modular water sampling systems that utilize hollow-membrane filtration cartridges were developed by RKS laboratories and tested by the Fisheries and Oceans, Canada, Molecular Genetics Laboratory. Compared to Sterivex filters, an industry standard for eDNA filtration, the hollow-membrane filtration cartridges allowed for a six-fold increase in filtration volume and threefold increase in filtration speed. The field sampling systems, which combine pumps, a programmable controller, an air pump, an ozone generator, and up to eight filters at once, enabled efficient direct eDNA filtration from diverse aquatic environments, from creeks to the open ocean. To evaluate ease of deployment, we present the results of a three day workshop where technical staff of an Indigenous resource management organization, without any prior knowledge in eDNA sampling, were trained and performed independent eDNA sample collection. The samples were analyzed by metabarcoding and qPCR to reveal the distributions of salmon and other species co-occurring in salmon ecosystems, from large ephemeral predators, to the planktonic prey of salmon, even including their pathogens. In this example study, we further observed a substantial shift in community composition in the vicinity of aquaculture facilities where marine species associated with aquaculture feed were detected in freshwater at high relative abundance. This study demonstrates how these sampling systems provide an efficient entry point for small and medium scale organizations to utilize eDNA to fulfill their research and monitoring objectives.

## Introduction

Environmental DNA (eDNA) utilizes traces of DNA in the environment to provide an account of the diversity and distribution of aquatic organisms, from microorganisms to whales, without the need for capturing or handling individuals (Rees et al. 2014). Applications like invasive species monitoring have been at the forefront of the eDNA revolution (Rees et al. 2014; Thomas et al. 2020; Westfall, Therriault, and Abbott 2022) but the approach is also gaining a foothold in ecology due to its ability to provide semi-quantitative insights into species composition comparable to, and often more sensitive than, conventional sampling methods (Jerde, Wilson, and Dressler 2019; Spear et al. 2021; Lacoursière-Roussel et al. 2016; Stoeckle et al. 2020; Thomsen et al. 2016; Deeg et al. 2023).

Our program carries out research on the health of salmon and their ecosystems, with particular focus on identification of factors contributing to the multi-decadal trend in declining salmon survival in the Pacific Northwest (Crozier et al. 2021; Wilson et al. 2022). Focusing on salmon infectious health (Bass et al. 2022, 2023; Deeg et al. 2022), predation (Miller et al. 2014; Furey et al. 2021), climate change and environmental stress (Akbarzadeh et al. 2020; Houde et al. 2019; Jeffries, Hinch, and Sierocinski 2014), salmon hatcheries (Nekouei et al. 2019), aquaculture (Shea et al. 2020; Bateman et al. 2022) and catch-release fisheries (Cook et al. 2015), we have applied molecular approaches to understand factors contributing to survival of salmon in their natural habitats. In recent years, our program has shifted to include eDNA assessments to ascertain the health of salmon ecosystems by characterizing co-distributions of salmon, their predators, prey, pathogens, and competitors, merged with holistic assessments of salmon health (Deeg et al. 2023, 2022). With specific interest in early marine survival of salmon, we recognized the need for an efficient, high volume filtration system for collection of eDNA samples in the ocean – from shore, small boats/drones, and large ships – that could enable collection of dozens of high volume (multi-litre) samples in a day.

eDNA studies in marine ecosystems lag those in freshwater, in part owing to the need for higher filtration volumes and sampling intensities to describe vast ocean ecosystems (McClenaghan et al. 2020; Govindarajan et al. 2022; Robinson et al. 2023). Traditionally eDNA has been collected on flat filters, which are inexpensive but difficult to handle without risking contamination in the field. Cartridge Sterivex filters, designed for sterilizing liquids, allow for easier handling and reduced contamination risk, but have lower filtration capacity, often in the 100’s of mls (0.22 um PES filters) to 1-2 L (0.45 um PVDF filters). Moreover, slow flow rates of Sterivex, specifically PVDF membranes, routinely require 30-60 minutes to obtain these volumes, forcing researchers to bring water samples back to the laboratory for filtration. This risks nucleic acid degradation and contamination and requires additional labor compared to systems capable of in-field filtration.

To circumvent these issues and make eDNA sampling accessible to a wider user base, different organizations and companies have attempted different solutions. The Japanese ANEMONE project (https://anemone.bio/) forgoes the need to transport water samples to a laboratory by filtering in the field using simple syringes equipped with Sterivex (EMD Millipore) cartridge filters (Suzuki-Ohno et al. 2023). This approach allows for samples to be collected with minimal equipment and minimal risk of contamination as the filtration membrane is protected in a cartridge which also allows for immediate sample preservation with RNALater or other preservatives. However, the low filtration capacity of Sterivex filters, especially when manually filtering with a syringe, limits the filtration volume with this approach to 500ml, thereby limiting sensitivity. On the other hand, Smith-Root (Vancouver, WA, USA), a fisheries equipment manufacturer, offers customized backpack samplers (Thomas et al. 2018). These sampling systems deploy flat filters with large nominal pore sizes (up to 5µm) to improve flow rate and filtration volumes and require careful handling of flat filters or specialized filter preservation systems.

First Nations Peoples in British Columbia have long-standing relationships with wild Pacific Salmon and prior to colonization managed sustainable salmon fisheries for at least 5,000 years (Campbell and Butler 2010; Atlas et al. 2021). While the abundance of numerous salmonid populations in BC has declined significantly due to anthropogenic impacts, salmon remain integral to the culture, food security, and wellbeing of Indigenous Peoples in BC and around the Pacific rim (Tabarev 2011; Muckle 2011; Yoshiyama 1999; Lichatowich and Lichatowich 2001). As rights and title holders, First Nations maintain an ancestral and ongoing relationship as stewards of the natural world, providing vital leadership towards the protection and rebuilding of wild salmon within their traditional territories and throughout the salmons’ migratory life cycle (Atlas et al. 2021; Reid et al. 2021). Many Indigenous-led salmon science initiatives could benefit from the application of eDNA tools to address key data gaps and inform management and conservation actions. However, substantial barriers to accessing this powerful technology remain for First Nations. Perceived complexity of sample collection, uncertainty about data interpretation, risk of contamination, and inaccessibility of analysis providers are amongst the top impediments.

Here we evaluate a simple eDNA sampling system developed by RKS-labs that relies on protected hollow-membrane cartridge filters and customized modular sampling systems that allow for highly efficient eDNA sampling. To demonstrate the feasibility of this sampling approach for salmon ecosystem monitoring, we present the results of a single day eDNA survey performed by Indigenous fisheries managers after two days of training.

## Materials and Methods

### Hollow-membrane filtration cartridges

The purpose-built filtration cartridges (RKS Laboratories LTD, French Creek, BC, Canada; “EZ-E-DNA”; patent pending # 18/313335) contain 120 Polyethersulfone (PES) hollow-membrane tubules with a nominal pore size of 0.45 µm that provide an increased active filtration surface area compared to flat filters or conventional cartridge filters (Figure 1). The compact cartridges are equipped with female luer-lock connections on both ports and can withstand up to 60 PSI of pressure.

**Figure 1:**
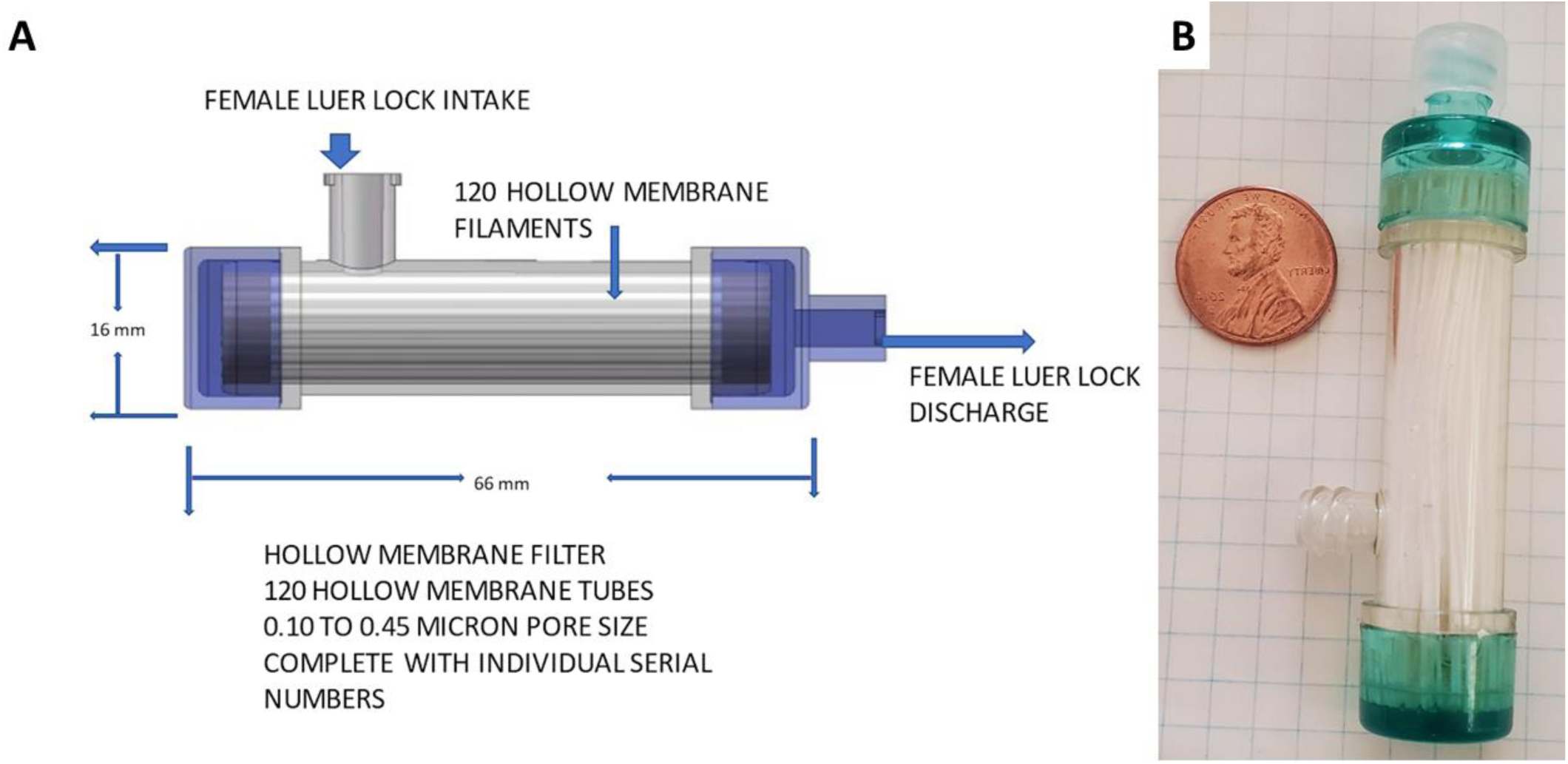
Hollow-membrane cartridge filters by RKS Labs: A Schematic B: Filter size compared to a US penny.

### Sampling systems

Sampling systems were designed to collect water and simultaneously filter it through cartridge filters (Figure 2; RKS Laboratories LTD, French Creek, BC, Canada; “EZ-E-DNA”; patent pending # 18/313335). Described in the following is the “standard model” system that was developed for deployment from small vessels (Figure 2A,B, Figure 3A,B). Water is collected at depth through an intake line that is attached to a downrigger equipped with a 10lb weight (model “Depth Power”, Scotty manufacturing LTD. Sidney, BC, Canada; Figure 3A,B). Inside a polymer case housing a 12V DC pump draws water through the intake line and delivers it to a combined flow counter and programmable controller unit (referred to as “controller” below). The controller monitors the flow rate and filtration volume and automatically stops filtration once the targeted volume is reached. The controller can be bypassed in order to prime the pump before filtration. An air pump is included to purge residual liquid from the system after filtration is completed and connects to the main system via a one way valve. From here, all lines lead to the master valve and consecutive valve controlled branches that allow for any combination between one and four replicate filters to be processed simultaneously or sequentially. The filtrate is discarded through waste lines. The standard sampler can also be deployed for collection from shore when the intake line is attached to an extendable aluminum pole so that the sample can be collected mid-water column away from any obstacles or sediment (Figure 3C). A backpack mounted version has also been developed and tested. In addition to the standard model, a dedicated sampling system to integrate into existing oceanographic sampling workflows that collect water at depth via Niskin bottles (Figure 2C,D, Figure 3D) has been developed. This system (“milk machine”) allows for simultaneous filtration from four Niskin bottles from different depths and has already been deployed on open ocean research expeditions to the NE Pacific (Figure 3D).

**Figure 2:**
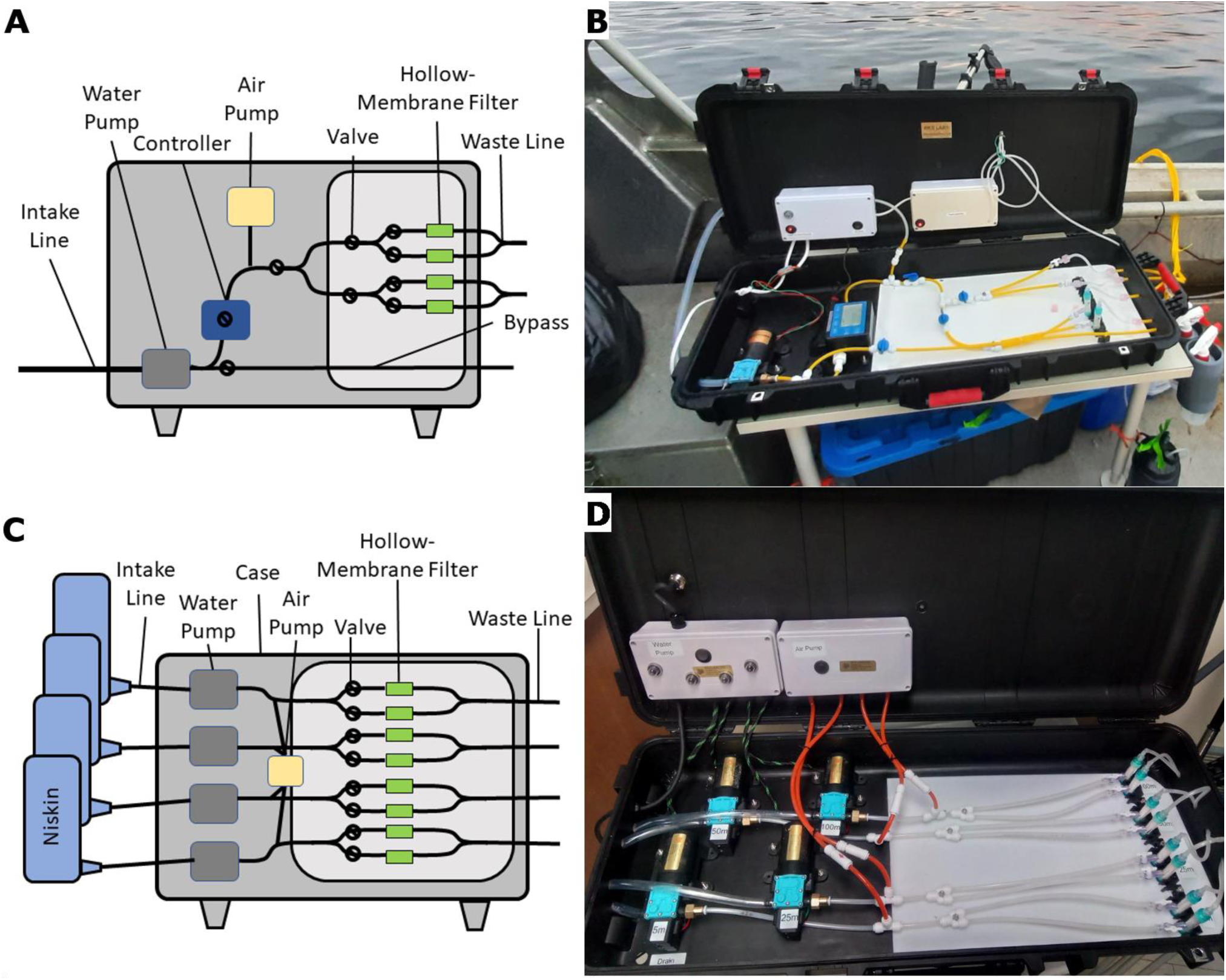
RKS EZ-eDNA sampling systems: A:Standard model schematic B: Standard model overview, F: “Milk machine” model schematic, G: “Milk machine” model overview.

**Figure 3.**
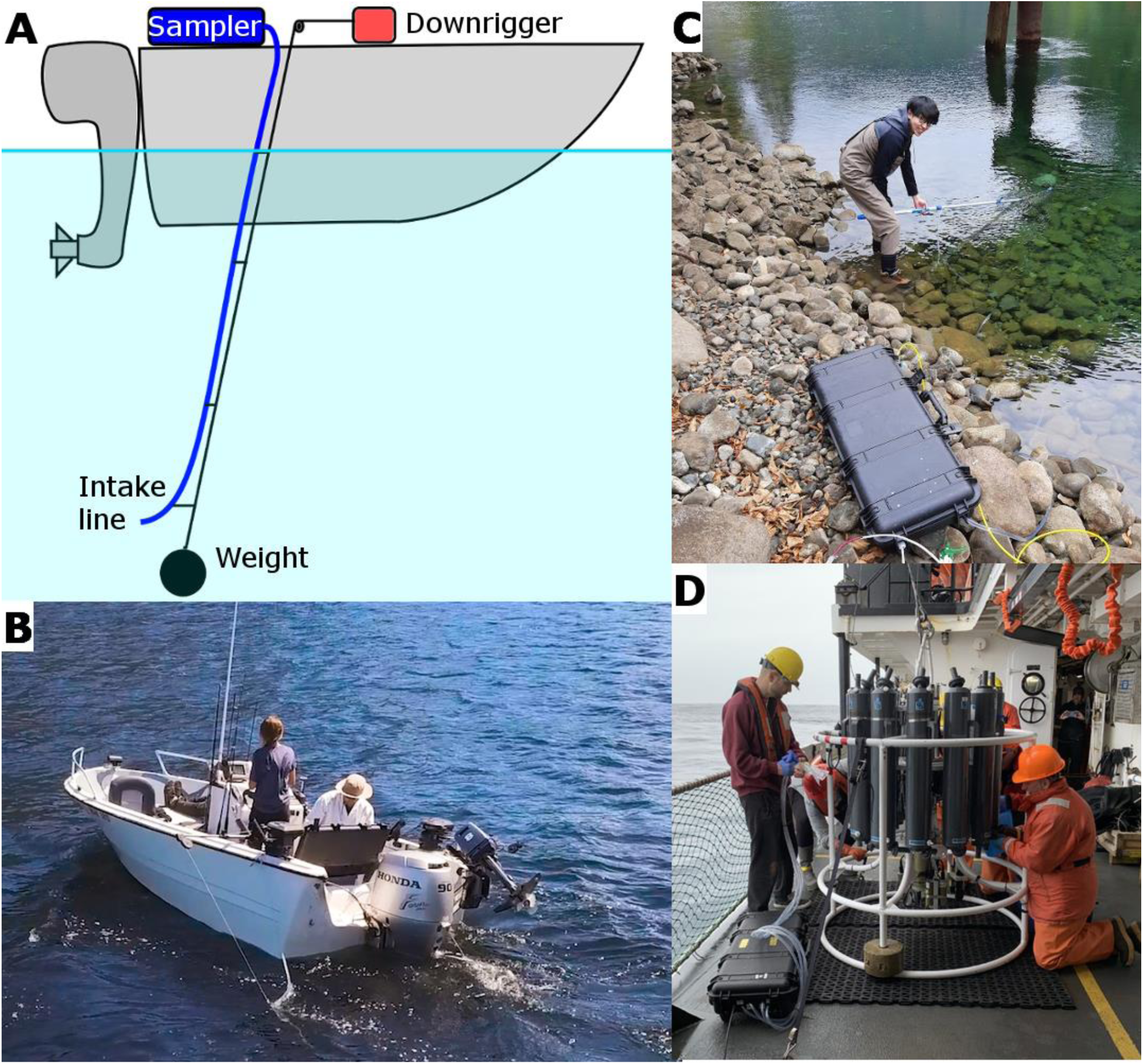
RKS EZ-eDNA sampling systems in the field A: Schematic of standard model (blue) deployment from small vessel via downrigger (red), B: The authors operating the standard model (blue) coastal deployment from small vessel via downrigger, C: The authors operating the standard model deployment in a small stream via pole D: The authors simultaneously sampling from four Niskin bottles from differing depths with the “Milk machine” model.

### Sample collection (standard model)

The standard sampling system is decontaminated in the laboratory before field deployment by recirculating 1.6% sodium hypochlorite through the system for 15 min, followed by a flush with ultrapure 2L MilliQ water (MQ: RO-water UV-treated at 18.2 MΩ·cm; EMD Millipore), followed by 5 min recirculation of 0.1 N sodium thiosulfate solution, and a final flush with 2L MQ water. Once the cleaning is completed the sampling kit is kept closed until collecting the first sample and the intake line is protected with a Whirl-Pak. Similarly, all surfaces in the sampling area onboard the vessel are thoroughly decontaminated by spraying and wiping with 1.6% sodium hypochlorite, followed by MQ water, 0.1 N sodium thiosulfate solution, and MQ water again. An ozone generator was run continuously to ozonate a 2L bottle of MQ water for decontaminating the sampling system between sampling sites. Consumables needed at each sampling site include a bench cover, two hollow-membrane filters, one 5ml syringe filled with RNA later, four male Luer stops, labels, and a Whirl-Pak.

At each sample site, the system is initially flushed with 20L of sample water. For this, a connector is inserted in place of the filters and all valves are opened to allow water from the sample site to pass through the lines without resistance and flush out any residual bleach or ozone. Next, the remaining liquid is purged from the system via the air pump and the appropriate number of hollow-membrane cartridge filters are inserted into the system. The controller is programmed to the desired filtration volume (usually 5L per filter) and filtration is started. Once the desired filtration volume has been collected the system automatically terminates the water collection and the remaining liquid in the hollow-membrane filtration cartridges is purged via the air pump. The hollow-membrane filters are recovered individually and injected with 2 ml of RNAlater via the intake Lure port before all ports are capped with Luer-Stop and collected in a Whirl-Pak. Samples are kept at 4°C and protected from direct sunlight. After sample collection, the system is set to recirculate 2L of ozonated MQ water (> 0.5 ppm O_3_) for 10 minutes without filters in place and all valves open to remove any remaining contaminants in the field.

For samples collected in the marine environment, the sampling hose is attached to a downrigger (model “Depth Power”, Scotty manufacturing LTD. Sidney, BC, Canada) and lowered to the desired sampling depth (up to 10m below surface). Both point and transect samples can be collected. To evenly represent transect samples, the water flow is throttled to approximately 0.2L / min / filter by partially closing and adjusting the valves to allow for a more constant filtration rate. For samples collected from shore in freshwater environments, the sampling hose is attached to an extendable 5m aluminum pole so that the sample intake was mid-water column away from any obstacles or sediment. Metadata such as Lat/Long, time, filtration volume, sampling depth, transect length, etc is recorded. To assess cross contamination between samples, “MQ water only” field control samples are collected before the first sample, after the last sample, as well as one sample mid day.

### Filtration capacity comparison

To assess the filtration capacity of Hollow-membrane (HM) filters and Sterivex filters (EMD Millipore, Burlington,MA, USA) of the same nominal pore size (0.45 µm) were compared head to head in the field. To compare the maximum filtration capacity, sediment rich water was collected 0.5m off the bottom in the Somass River in BC, Canada (16/02/ 2024; Figure 4 “Max_Volume”) and alternatively pumped through Sterivex or HM filters (n=8). When the flow rate dropped below 10mL/min filtration was terminated and the filtered volume recorded. To compare filtration speed, time to filter 1 L of seawater from the Strait of Georgia at a dock in Nanaimo BC (10/04/2024; Figure 4 “Speed_rep”) at 6 m below surface was recorded (n=10). Similarly, filtration speed was also compared from twelve locations in Barkley Sound (24/04/2024; Figure 4 “Speed”).

**Figure 4:**
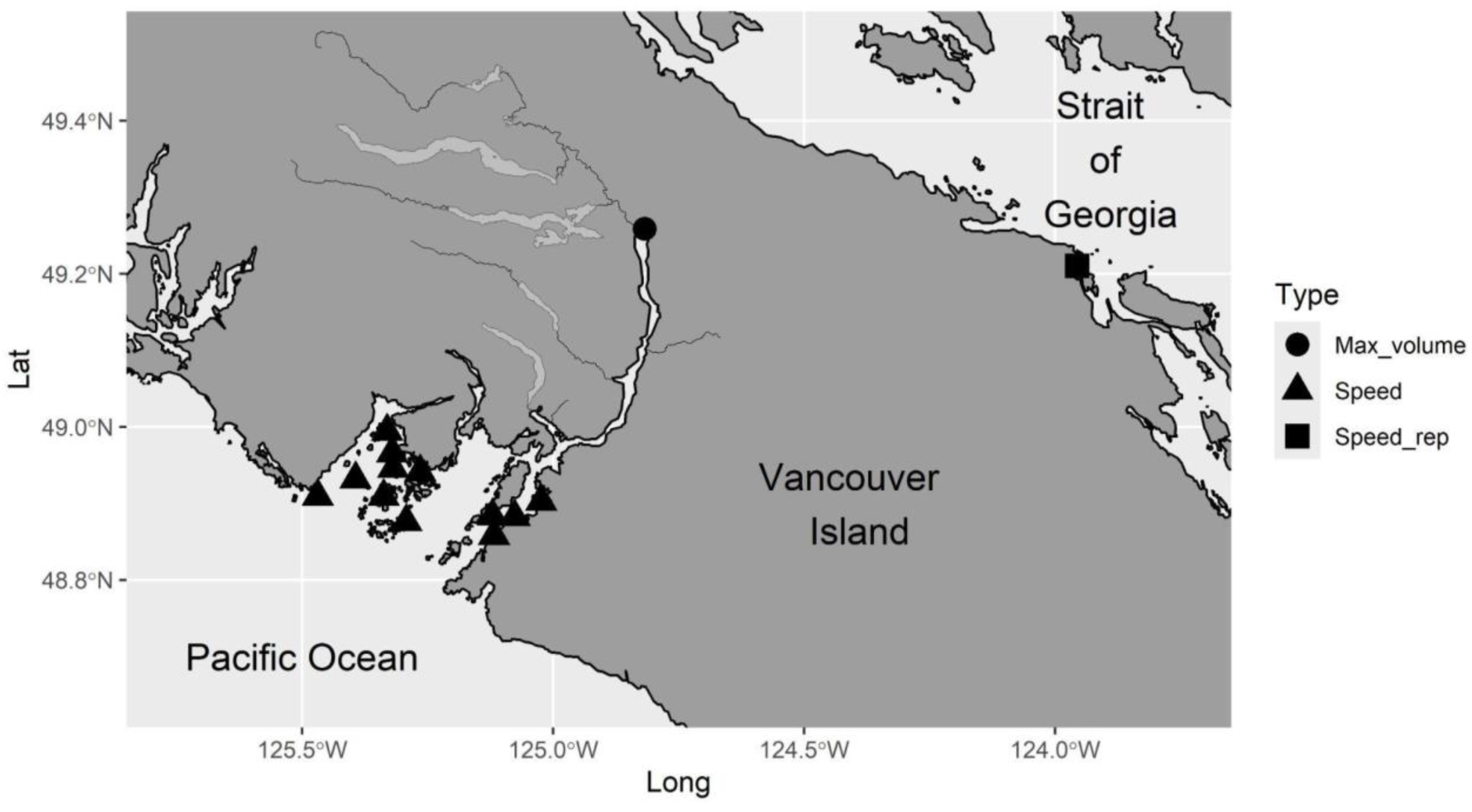
Sampling sites to compare filtration capacities. Samples were collected to compare maximal filtration volume (“Max_volume”), filtration speed with one sample per site (“Speed”), or filtration speed with multiple samples per site (“Speed_rep”)

### Uu-a-thluk Fisheries workshop and sampling

An eDNA sampling workshop was held Nov 29th to Dec 1st, 2022 in Port Alberni, BC for Uu-a-thluk Fisheries of the Nuu-chah-nulth Tribal Council. The goal of the workshop was to train participants in the use of eDNA as a non-invasive tool to assess species distributions with a focus on salmon as well as to assess the impact of industrial activities on ecosystems (e.g. accumulation and amplification of salmon pathogens or shifts in community composition). On the first day of the workshop participants were introduced to applications of eDNA and the underlying principles. On the morning of the second day sampling techniques were introduced, followed by an afternoon of hands-on practice with the RKS EZ-eDNA standard sampler. On the third day participants independently collected eDNA samples from a number of different environments of their choosing in the territories of the Huu-ay-aht, Tseshaht, Uchucklesaht, Yuułuʔiłʔatḥ, and Hupacasath nations. A total of 22 samples from 11 locations were collected during the workshop (Figure 5). One group of workshop participants set out by vehicle and foot to sample the Stamp River (Hatchery Lagoon), Robertson Creek, and Sproat River (Figure 5C). An additional 5 samples from the Stamp/Somass system were collected two weeks prior to the workshop (Figure 5C). The remaining samples were collected during the workshop from the Alberni Inlet and the Broken Island Group using the standard sampling system from small vessels via downrigger (Figure 5D). A total of four field-based control samples were collected by participants in the workshop which was well below the recommended control numbers and our internal standard sampling procedures..

**Figure 5:**
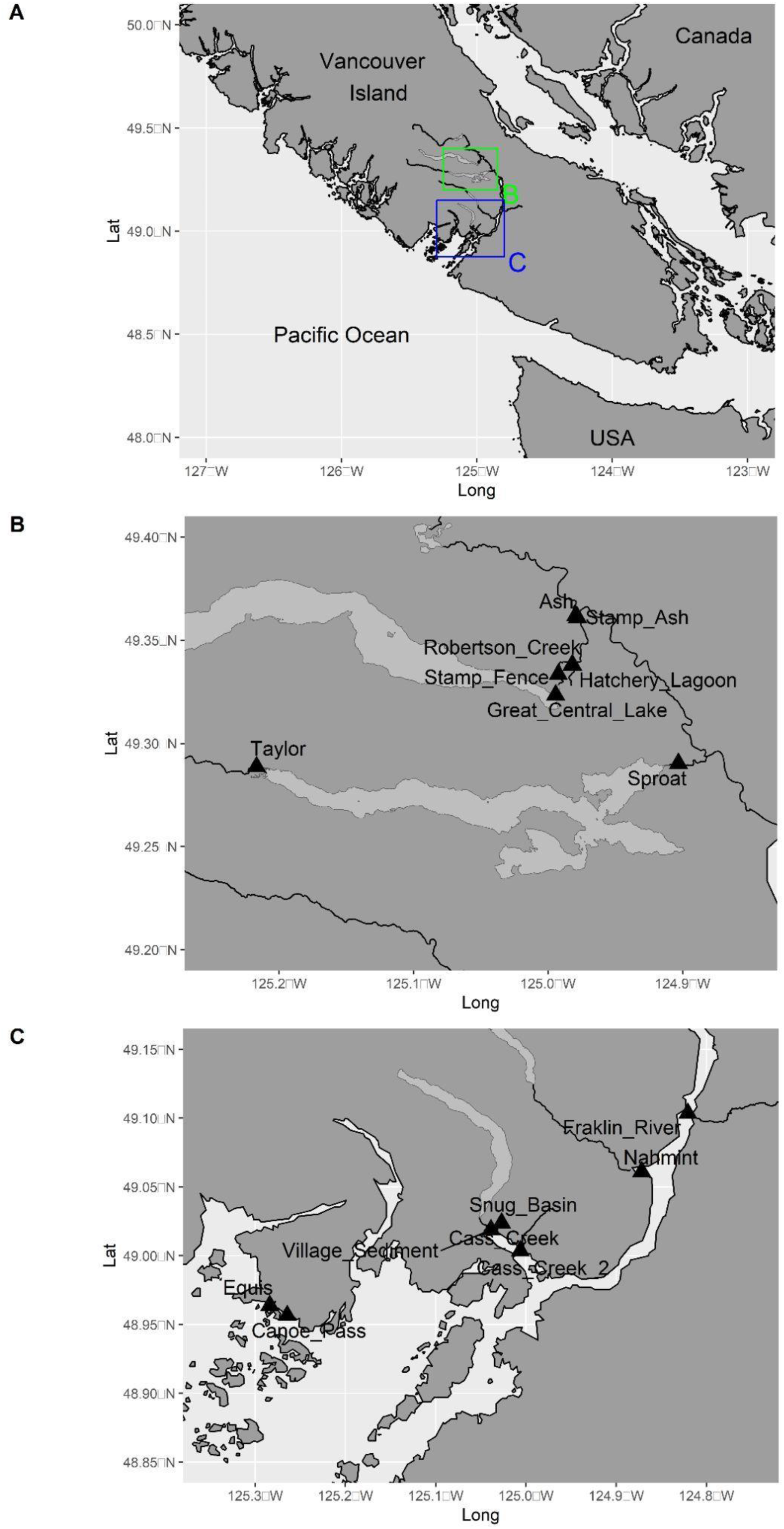
Uu-a-thluk fisheries workshop sampling sites: A: Location of sampling sites in Vancouver Island with highlighting the locations of inserts B and C. B: Somass/Stamp watershed. Sampling sites Hatchery lagoon and Robertson creek and Stamp_Ash and Ash are in close proximity and overlap partially in this figure.C: Alberni Inlet and and Broken island sampling sites. The two Cass Creek samples are in close proximity and overlap partially in this figure.

### DNA Extraction

Upon return from the field, DNA was extracted from the filter cartridges by purging remaining RNAlater by flushing the filter with 3 ml of purelink viral RNA/DNA extraction buffer (Invitrogen, Waltham, MA, US). Next, 2 ml of extraction buffer with proteinase K (20mg/ml; Invitrogen) was injected into the filter cartridge and incubated at 56℃ for 30 minutes while being agitated on a shaking platform at 1000 RPM for 10 min. After incubation, the lysate was removed from the hollow-membrane filter by backflushing it with 1 ml extraction buffer and centrifuged at 2400xg for 5 min. The supernatant was mixed with EtOH to a final concentration of 37%, vortexed, and transferred to a Zymo-Spin IIC spin column (Zymo) attached to a vacuum manifold. The columns were washed twice with 500 ul of Wash Buffer (WII, Invitrogen), dried by spinning at maximum rpm in a centrifuge and eluted with 30 ul of sterile RNAse-free water. Extraction yields were assessed using a fluorometer with the Quant-iT™ dsDNA HS kit (Thermo Fisher). Newly opened unused hollow-membrane filters were included in the extraction protocol as extraction blank controls to assess any contamination introduced by the extraction.

### Metabarcoding

Metabarcoding analysis was performed as described previously (Deeg et al. 2023). In brief, four amplicons targeting 16S for Chordates & Cephalopods (Deagle et al. 2009), 12S for Fish (Miya et al., 2015), and COI for Salmonids (Thomas et al. 2017) were amplified and libraries were prepared using the Kappa LT library prep kit (Kappa Bioscience AS) according to manufacturer’s instructions. Libraries were sequenced in a single Miseq SE 300 cycle run that was shared with thirteen additional samples from an unrelated project. Sequencing data was quality controlled, demultiplexed, filtered, and assigned to individual samples using OBItools (https://git.metabarcoding.org/obitools/obitools/wikis/home). Local BLASTn (https://blast.ncbi.nlm.nih.gov/Blast.cgi) was used against the NCBI nr database (version Jan. 30. 2023). MEGAN (https://bio.tools/megan) was used to assign BLAST results to the lowest taxonomic level. R version 4.0.3.was used for statistical analysis and visualization of the data. Reads belonging to contaminants such as humans and common terrestrial food species (such as pigs, chicken, and cows) were excluded from the analysis. Species detections with less than 10 reads per sample were disregarded (i.e. set to zero). For relative abundance comparison, an eDNA index was calculated for every species at every site that is equivalent to a Wisconsin double standardization (Kelly, Shelton, and Gallego 2019; Port et al. 2016).

### High-throughput quantitative polymerase chain reaction

Samples were also assayed for the presence of infectious agents of salmon as well as selected fish species (Supplementary Table 1) using qPCR on the the Fluidigm BioMark™ platform (Fluidigm Corporation, CA, USA), a nanofluidic automated real-time PCR system as described previously (Miller et al. 2016; Bass et al. 2017; Shea et al. 2022; Bass et al. 2017). In brief, following extraction, nucleic acids (RNA and DNA) were reverse-transcribed to cDNA. Next, cDNA/DNA underwent a pre-amplification step (17 cycles) using a 1/10 dilution of all primers (no probes) targeting sequences to be assayed on the BioMark (Miller et al. 2016). Following pre-amplification, samples and assays were analyzed on a 96 x 96 well dynamic array (Fluidigm Corporation) as per Miller et al. (2016). The data was analyzed for cycle threshold using the Fluidigm Real-Time PCR Analysis software. Potential laboratory contamination was assessed by internal artificial positive controls (APCs) and samples with APC detections were discarded.

### Controls

After removing human reads and common food associated contaminants, which can result from contamination by the collector and/or contamination in the watershed, the laboratory extraction control and three of the four field controls contained no remaining reads (Table 2). One remaining field control showed contamination from Pacific salmon (*Oncorhynchus*), Chinook salmon (*Oncorhynchus tshawytscha*) and water fleas (*Evadne nordmanni*). As this was a replicate of a control filter that showed no remaining reads (Field control 1a vs 1b) the contamination is presumably related to filter handling and no correction was made to the remaining dataset.

**Table 1:**
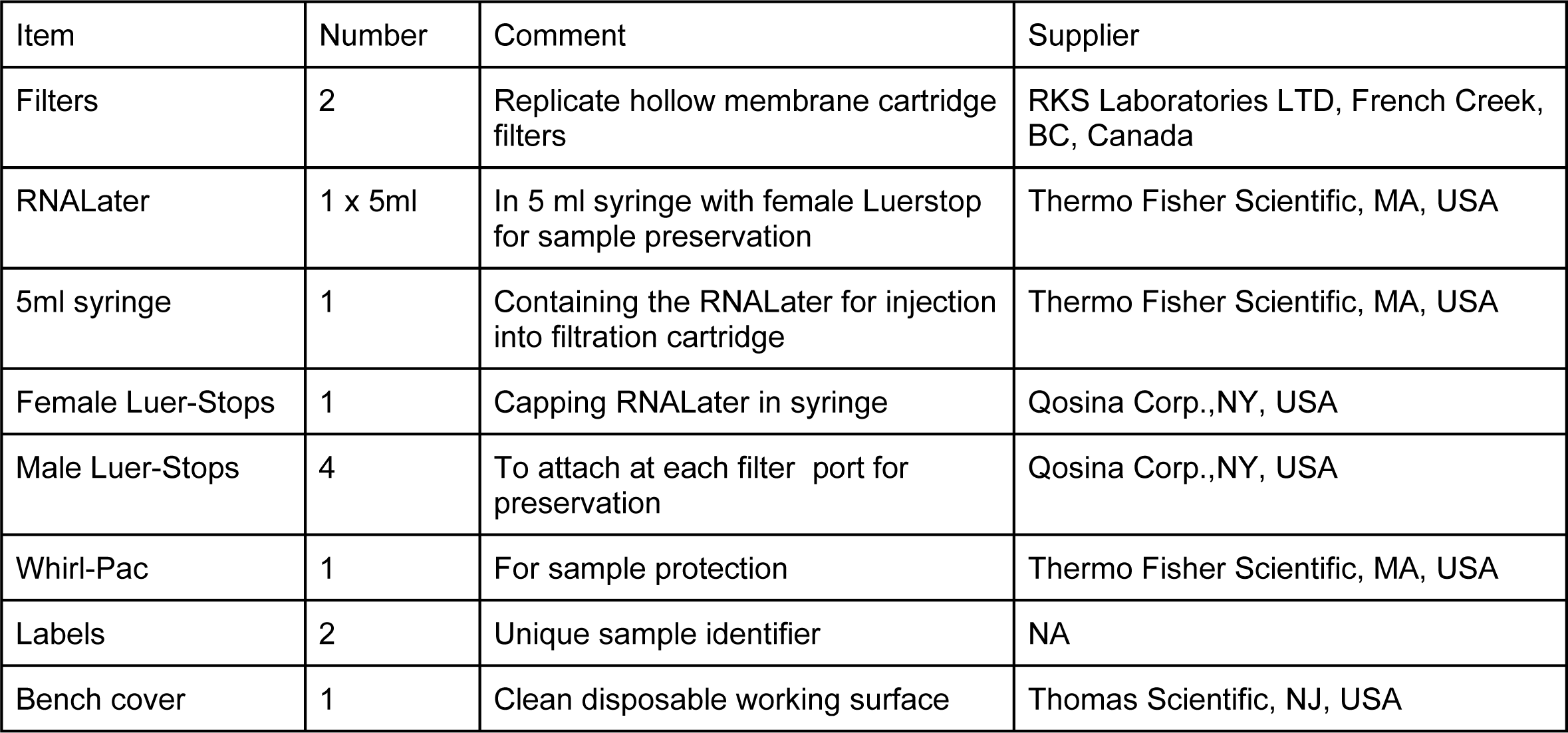
Site kit contents used in conjunction with the RKS EZ-DNA sampling systems.

| Item | Number | Comment | Supplier |
| --- | --- | --- | --- |
| Filters | 2 | Replicate hollow membrane cartridge filters | RKS Laboratories LTD, French Creek, BC, Canada |
| RNALater | 1 x 5ml | In 5 ml syringe with female Luerstop for sample preservation | Thermo Fisher Scientific, MA, USA |
| 5ml syringe | 1 | Containing the RNALater for injection into filtration cartridge | Thermo Fisher Scientific, MA, USA |
| Female Luer-Stops | 1 | Capping RNALater in syringe | Qosina Corp.,NY, USA |
| Male Luer-Stops | 4 | To attach at each filter port for preservation | Qosina Corp.,NY, USA |
| Whirl-Pac | 1 | For sample protection | Thermo Fisher Scientific, MA, USA |
| Labels | 2 | Unique sample identifier | NA |
| Bench cover | 1 | Clean disposable working surface | Thomas Scientific, NJ, USA |

**Table 2:**
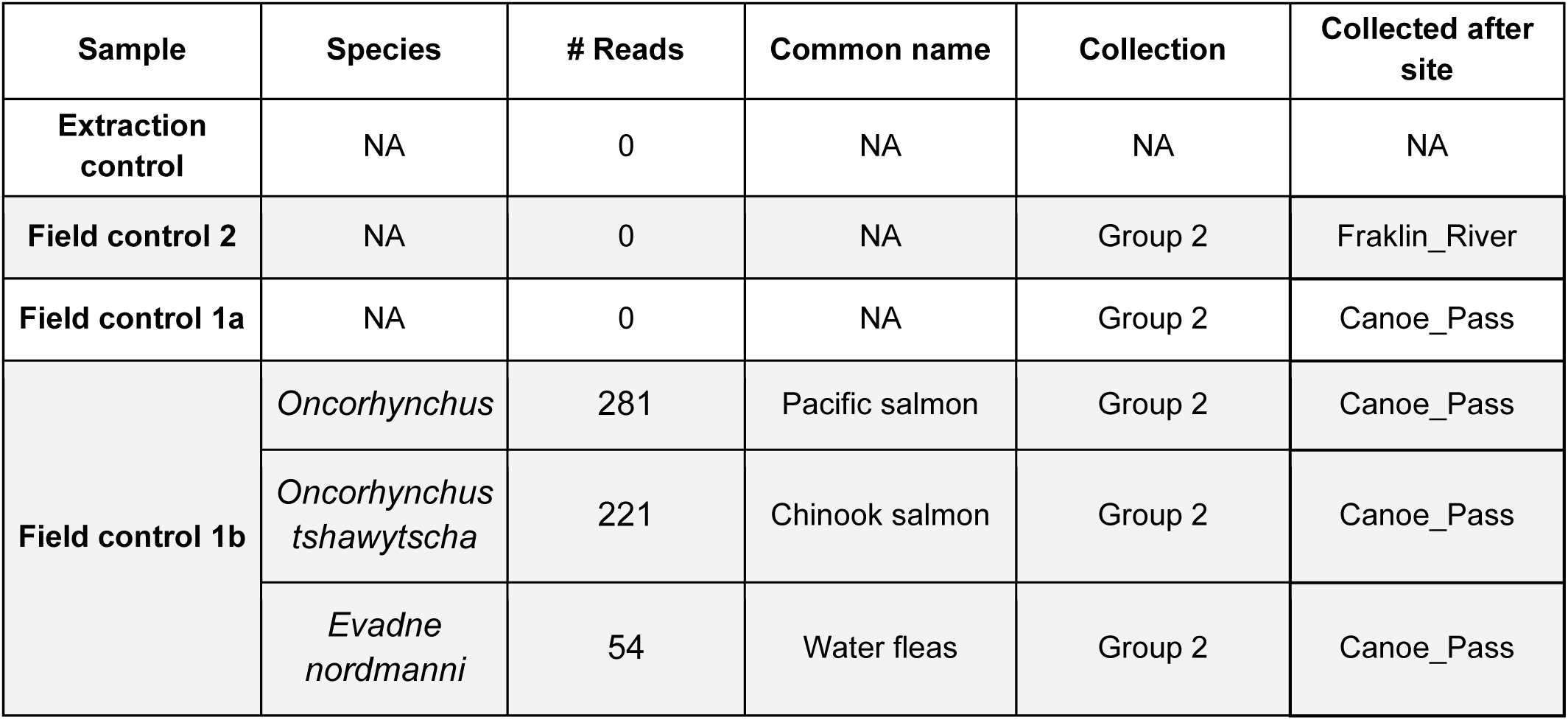
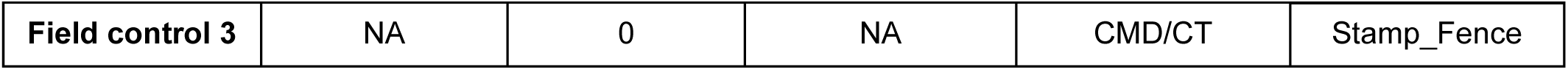
Reads and read numbers in control samples (after removing reads for non-target contamination species)

## Results

### Filtration capacity comparison

When comparing HM filters with Sterivex filters of matching filtration material and pore size (Sterivex PES 0.45 µm; HM nominal pore size 0.45µm), HM filters allowed for an approximately six fold increase in maximum filtration volume (HM: 3.64 ± 1.17 L; Sterivex: 0.61 ± 0.08 L; Figure 6A) in the test freshwater ecosystem with high sedimentation, although they showed a larger variability in filtration volume. Similarly, when filtration speeds were repeatedly compared head to head at a marine location in BC, HM filters were approximately three times faster than Sterivex filters in filtering 1L of seawater (HM: 33.1 ± 3.5 s; Sterivex: 114.0 ± 36.1 s; n=10; Figure 6B). The same speed advantage was confirmed when filtration speeds were compared head to head from samples collected at 8 m at twelve diverse marine locations across Barkley Sound in BC (HM: 53.9 ± 3.3 s; Sterivex: 135.6 ± 19.5 s; n=12; Figure 6C). For all of the Barkley Sound locations HM filters successfully filtered 5 L of water. Similarly, during the workshop, participants were successfully able to collect 5 L from all marine sites, but in the freshwater environment the mean filtration volume was 4.50 +/− 0.85 L.

**Figure 6:**
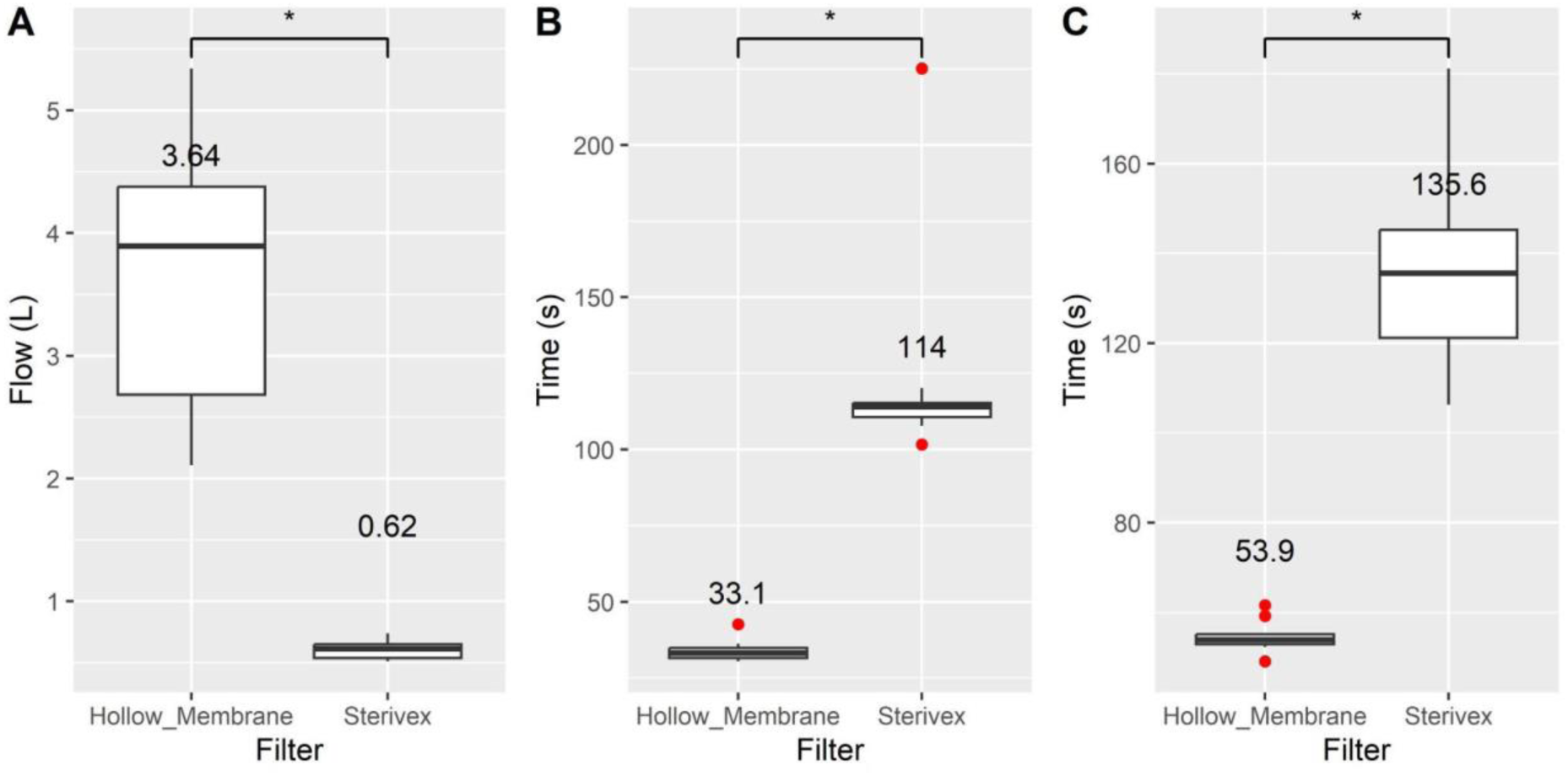
Pairwise filtration comparison between Hollow membrane and Sterivex filtration cartridges of matching poresize (PES 0.45 µm) A: Maximum filtration volume capacity in a sediment rich freshwater environment: Mean filtration volume (L) indicated above box plot. Two-sided student’s t-test p: 1.25 2.91×10^−5^ n=8. Note: two potentially defective HM filters–one that clogged at only 0.52L and the other that filtered 19.2L– were removed. B: Time to filter 1 L of seawater from the same location in Departure Bay BC: Mean filtration time (s) indicated above box plot. Two-sided student’s t-test p: 2.91×10^−5^ 0.0000291. n=10. C: Time to filter 1 L of seawater from twelve matched marine locations across Barkley Sound BC. Mean filtration time (s) indicated above box plot. Two-sided student’s t-test p: 1.01×10^−8^. n=12.

### Uu-a-thluk workshop metabarcoding results

#### Freshwater: Somass/Stamp watershed

Environmental DNA samples were collected from across the Somass/Stamp watershed. The Stamp mainstem was sampled from upstream to downstream immediately below Great Central Lake, downstream at the “fence”, at a lake like section at hatchery lagoon and immediately before the confluence with the Ash River (“Stamp at Ash”). Several tributaries of the Stamp River were also sampled, namely Robertson Creek, Ash River, and Taylor River that is a tributary of Sproat lake as well as Sproat River immediately below the eponymous lake (Figure 5). The “Stamp at Fence’’ site is approximately 500 m downstream of the discharge of a Cermaq hatchery (“Boot Lagoon Hatchery”) rearing Atlantic salmon (*Salmo salar*) for aquaculture and the site “Great Central Lake” is immediately upstream of the discharge from this facility. Similarly, the Robertson Creek Salmon Enhancement Program (SEP) facility run by Canada’s federal Department of Fisheries and Oceans (DFO) is rearing Pacific salmon (*Oncorhynchus spp.*) and discharges in the eponymous creek before it flows into the Stamp at Hatchery Lagoon.

Below Great Central Lake, the Stamp exhibited low species diversity, with Chinook (*O. tshawytscha*), coho (*O. kitsutch*), and sticklebacks (Gasterosteus. spp), as well as water fleas (*Eubosmina sp*.) and sculpins (*Cottidae*) being the primary species detected (Figure 7; Supplementary Figure 1, Supplementary Table 2). Downstream of the Cermaq facility (“Stamp Fence”), the native fauna, consisting of Chinook, coho, sticklebacks, and cutthroat trout (*O. clarkii*) was interspersed with reads assigned to Atlantic salmon (*S. salar*) as well as a number of marine fish species from the Atlantic and Pacific and Indian Oceans (Figure 7; Supplementary Figure 1, Supplementary Table 2). We presume that these detections of marine fish emanate from the feed given to hatchery reared salmonids. Similarly, at Hatchery Lagoon the sample collected near the sediment was extraordinarily rich within marine species, again presumably introduced to the system through hatchery feed. Native species were coho, stickleback, Chinook, and northern lampreys (*Petromyzontidae*) next to abundant invertebrates taking advantage of the nutrient rich environment. Robertson Creek itself, draining the eponymous SEP hatchery, was dominated by Chinook and coho reads, as well as aquaculture feed-associated taxa. Also detected were predators like black bear (*Ursus americanus*) and invasive species like bullfrog (*Lithobates catesbeianus*) and pumpkinseed (*L. gibbosus*). Immediately before its confluence with the Ash River, aquaculture associated reads were still detectable in the Stamp. The native community was rich in Chinook, sticklebacks, coho and invertebrates, with lower read numbers of black bears, bald eagles (*H. leucocephalus*), and northern lampreys.

**Figure 7:**
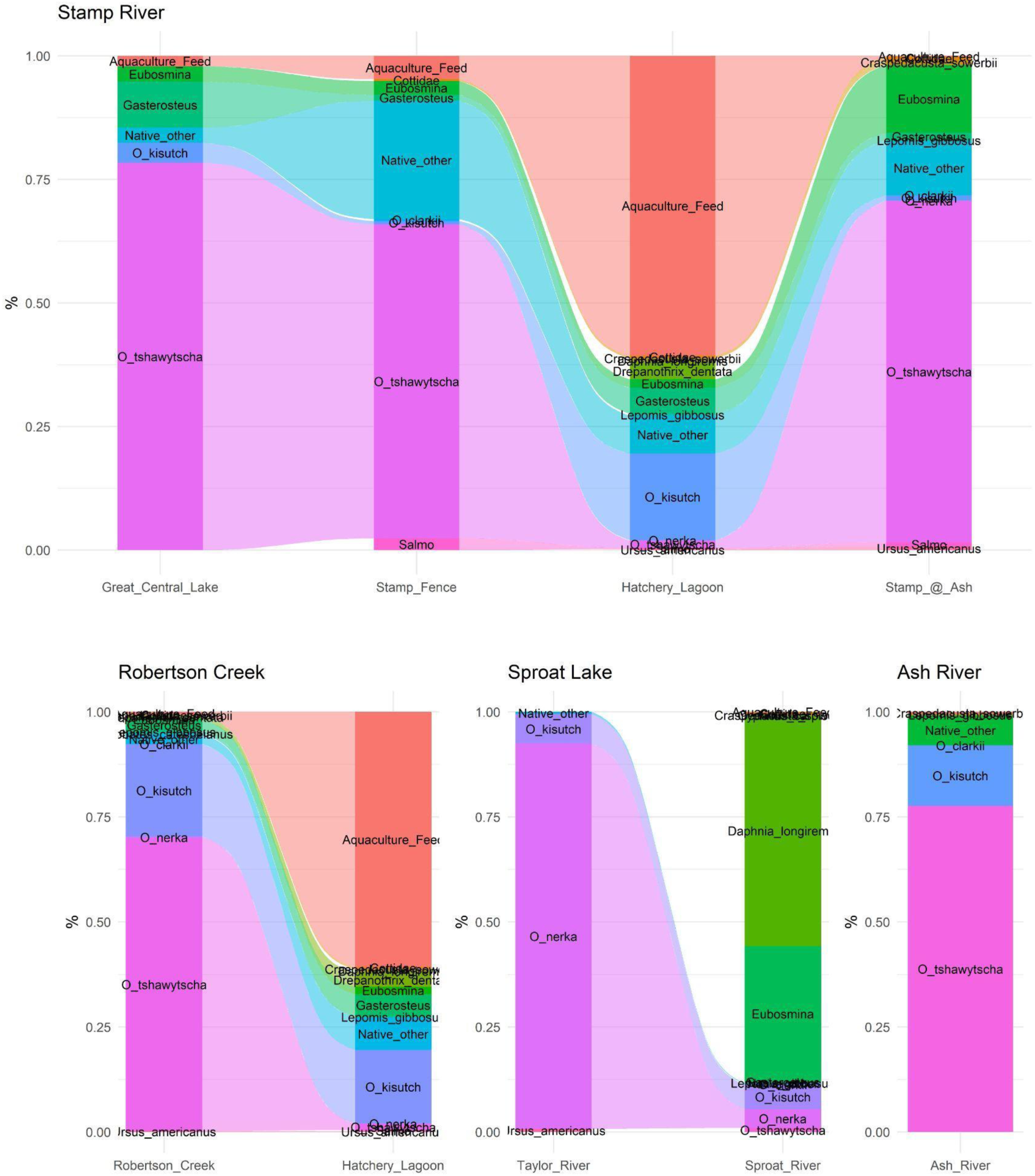
Community composition in the Somass/Stamp watershed based on metabarcoding data. High abundance species only (reads >450), genus level salmonid assignments removed (*Onchorynchus*). Marine species summarized under “Aquaculture_Feed’’.

The Taylor River was the least diverse site, dominated by sockeye (*Oncorhynchus nerka*) and coho, with a small number of black bear reads. Downstream of Sproat Lake, Sproat River was more diverse with abundant water fleas (*Daphnia longiremis* and *Eubosmina sp*.), coho, sockeye, and Chinook. Also detected were common carp (*Cyprinus carpio*) and pumpkinseed, two known invasive fish species. On the other side of the watershed, the Ash River also presented a low species diversity, consisting of abundant Chinook and coho and fewer cutthroat trout reads as well as invertebrates and predators of salmon.

#### Saltwater: Alberni Inlet / Broken Islands Group

Outside the Alberni Inlet, samples were uniformly dominated by reads assigned to the crustacean water flea *Evadne nordmanni* (Supplementary Table 3). The remaining species compositions differentiated between salmonid dominated river estuary communities and bivalve dominated coastal communities (Figure 8, Supplementary Figure 2-3, Supplementary Table 3). The Nahmint River estuary displayed the lowest number of reads of any marine sample and was rich in bay mussels (*Mytilus trossulus*), coho, and chum (*O. keta*). Marine forage fish like herring (*Clupeoidei*), stickleback, and anchovies (*Engraulis spp.*) were also detected alongside seals (*Phocidae*). The Franklin River estuary showed high read numbers of coho and Chinook, followed by chum and sockeye. Bay mussels were again abundant alongside sculpin, stickleback, barnacles and other invertebrates, and seals. The two samples near Cass Creek were dominated by water flea (*E. nordmanni),* followed by bivalves like bay mussel, saltwater clams (*Kellia*), the invasive milky Pacific venus clam (*Compsomyax subdiaphana*), forage fish like herring, surfperch (*Embiotocidae*), shiner perch (*Cymatogaster aggregata*), anchovy, and several sculpins, Pacific salmon, benthic invertebrates including sea stars (*Asteroidea spp.*), and rockfish (*Sebastes spp.*). The sample collected near the village sediment showed a similar composition with the exception of water fleas (*Eubosmina*) being highly abundant and the additional detections of California Sea Lions (*Zalophus*). Snug basin was also dominated by water fleas (*E. nordmanni*), followed by kelly clams and bay mussels. The fish community here was rich in herring, surfperch (*Embiotocidae spp.*), stickleback, Chinook and chum salmon. Hydrozoans (*Obelia*), ribbon worms, and ochre sea stars (*Pisaster ochraceus*) were the dominant invertebrates. The two most marine samples, Canoe Pass and Equis, were again dominated by *E. nordmanni,* but otherwise presented unique communities: The community at Equis was diverse with high read numbers of Pacific geoduck (*P. generosa*), benthic invertebrates, including nemertean worms (*Tubulanus sp.*), ochre sea star (*P. ochraceus*), leather star (*Dermasterias imbricata*), and bat star (*Patiria miniata*), tube-building spionid polychaetes (*Pseudopolydora paucibranchiata*), sea snail (*Petaloconchus montereyensis*), mossy chiton (Mopalia), nemertean worms (*Tubulanus spp.*), horseshoe worms (*Phoronopsis harmeri*), octopus (*Octopodidae spp.*), and sea cucumber (*Apostichopus parvimensis*). A diversity of fish were also detected, including Pacific herring, padded sculpin (*Artedius fenestralis*), kelp greenling (*Hexagrammos decagrammus*), coho, gunnel (*Pholis spp.*), sardines (*Sardinops spp.*), eelpout (*Lycodes spp.*), goby (*Coryphopterus spp.*), pricklebacks (*Stichaeidae spp.*), chimeras (*Hydrolagus spp.*) and rockfish (*Sebastes spp.*). The sample from Canoe Pass presented marine invertebrates including copepods, water fleas (*E. nordmanni*), ribbon worms and sea stars, most prominently the ochre sea star (*P. ochraceus*), Octopus, sea snails, crinoids (*Antedonidae spp.*), polychaetes (*Pseudopolydora spp.*), sea stars, red sea urchin (*Mesocentrotus franciscanus*) and several barnacles. Bivalves included bay mussels, Pacific geoduck, and the milky pacific venus clam (*C. subdiaphana*). Forage fish were also abundant with herring, greenlings, pricklebacks, kelp perch (*Brachyistius frenatus*), and stickleback present. Coho, humpback whales (*Megaptera novaeangliae*), and seals concluded this diverse community.

**Figure 8:**
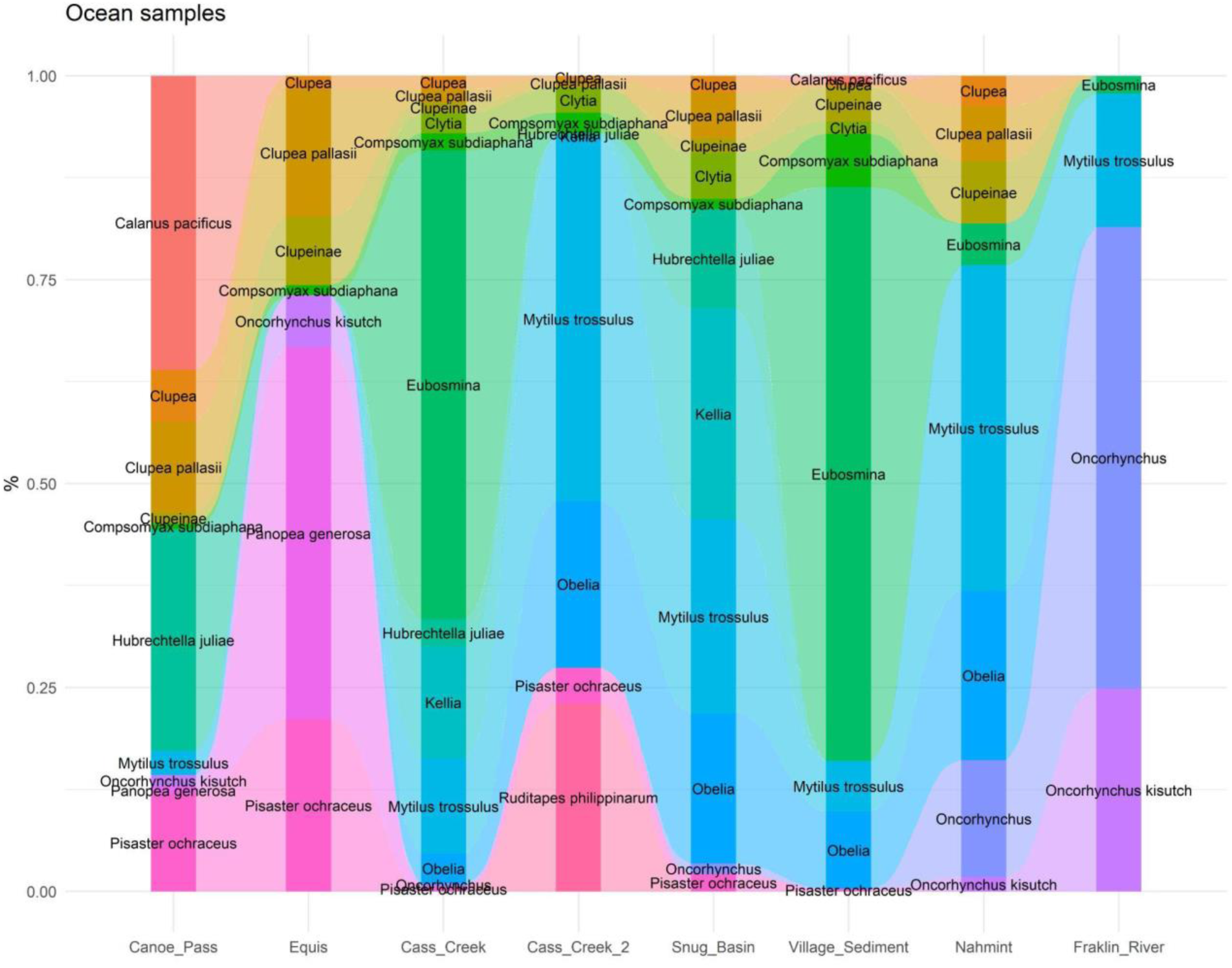
Community composition in then Alberni Inlet / Broken Islands Group based on metabarcoding data. High abundance species only (reads >450). Samples are arranged from marine to estuary influenced from left to right.

### Uu-a-thluk workshop quantitative PCR results

To demonstrate how eDNA data can be assessed with a targeted approach, we interrogated a subset of samples using qPCR assays for salmon and forage fish species (Figure 9). Pacific salmon species were detected at the highest concentrations in the river environment (e.g. sockeye at the Taylor River) and downstream of the SEP facility housing Pacific salmon (e.g. Chinook at Robertson creek; Figure 9). Similarly, Atlantic salmon were only detected in freshwater downstream of the Cermaq facility at Boot Lagoon (sites Robertson Creek and Hatchery Lagoon). Samples from the marine environment generally showed few salmon detections (over winter) and the existing detections were associated with freshwater influence such as the estuaries of the Nahmint and Franklin Rivers (Figure 9). Similar to the metabarcoding data, the qPCR data also showed very high detections of herring and anchovies in freshwater, specifically near salmon hatcheries, that were higher than marine detections and presumably are derived from aquaculture feed (Figure 9). The low level detection of anchovies in the Sproat is presumably associated with contamination (sample collected after Hatchery Lagoon). Marine detections of anchovies and herring were made at Cass creek, Snug basin (only anchovy), Equis, and Canoe Pass (only herring). The detection of Atlantic salmon in the marine environment at Snug basin is surprising and could originate from farm escapes or contamination. Clustering of estuarine samples (Nahmint, Franklin River) with true freshwater samples (Sproat, Taylor) shows the strong influence of river runoff in these estuarine samples. Aquaculture associated staples (Hatchery Lagoon and Robertson Creek) formed their own cluster due to the abundant feed associated detections (Figure 9)

**Figure 9:**
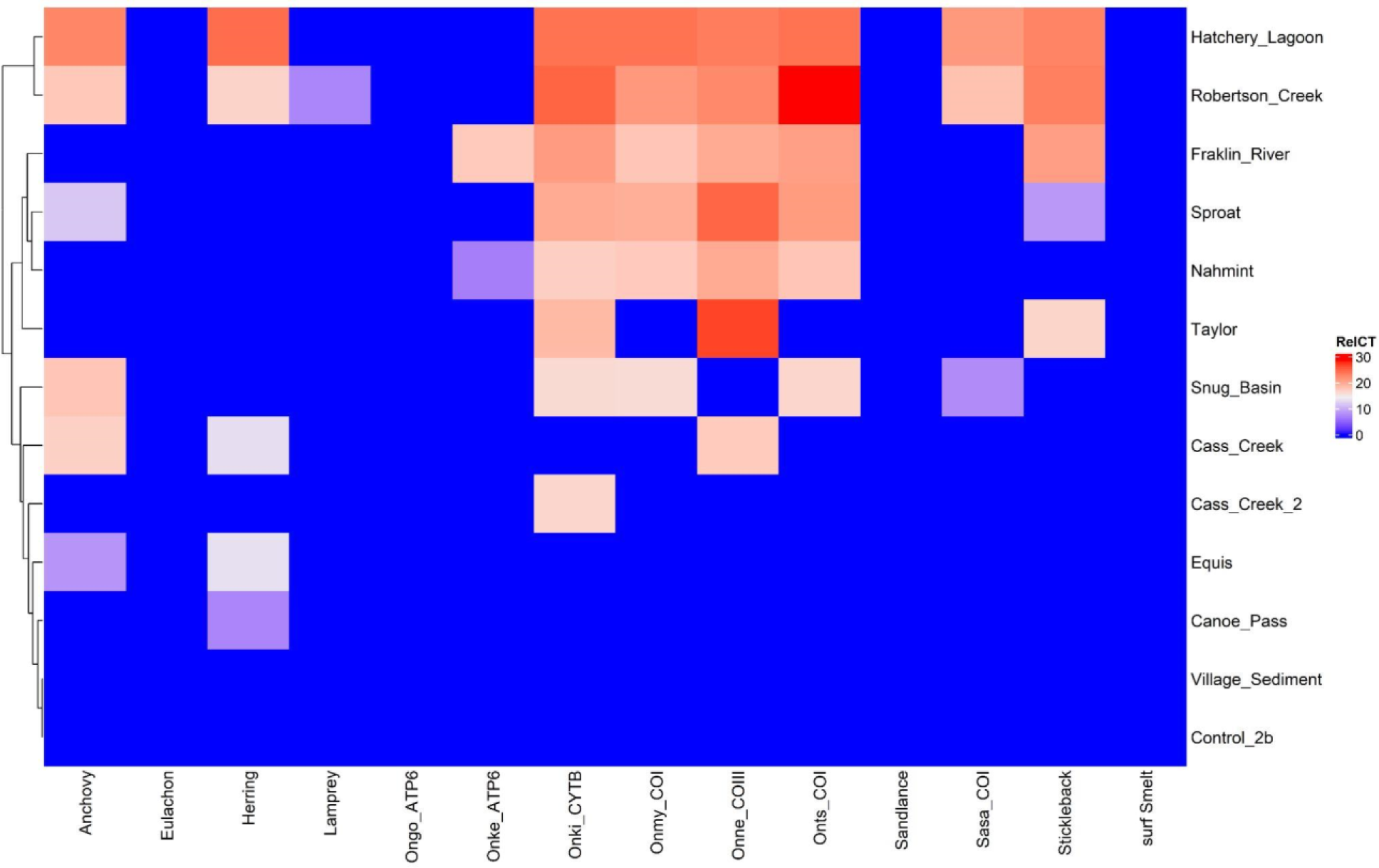
qPCR detections of fish species, RelCT depicted (45-CT). See Supplementary Table 1 for qPCR assay details. The sites at Great Central Lake, Ash river, Stamp at Fence, and Stamp at Ash were not tested for fish by qPCR.

Similar to screening for fish species, qPCR can also be utilized to screen for infectious agents of salmon. Accordingly, we assayed all samples for the presence of 31 infectious agents of salmon (Figure 10). The freshwater ciliate *Ichthyophthirius multifiliis* and the bacterium *Flavobacterium psychrophilum* were detected in all samples (the latter was also detected at low abundance in controls). The gill bacterium *Candidatus* Branchiomonas cysticola was only found in freshwater and estuaries, while the intracellular gill bacterium *Candidatus* Syngnamydia salmonis was only detected in marine samples (as well as in both controls). *Ichthyophonus hoferi*, a common marine parasite with a broad host range, was detected in most marine samples. In contrast, the ectoparasites *Ichthyobodo spp* were primarily detected in freshwater. *Loma salmonae*, a parasite often found in adult salmon, was accordingly primarily found where spawning adults were abundant. *Renibacterium salmoninarum*, a common bacterium found in aquaculture and returning adult salmon was accordingly found only downstream of the Cermaq and SEP aquaculture facilities, as was *Yersinia ruckeri*. The virus, SPAV-1, which primarily infects Chinook, was associated with detections of that species. A parasite of diverse marine fish species, *Kudoa thyrsites,* was only detected downstream of the Cermaq facility.

**Figure 10:**
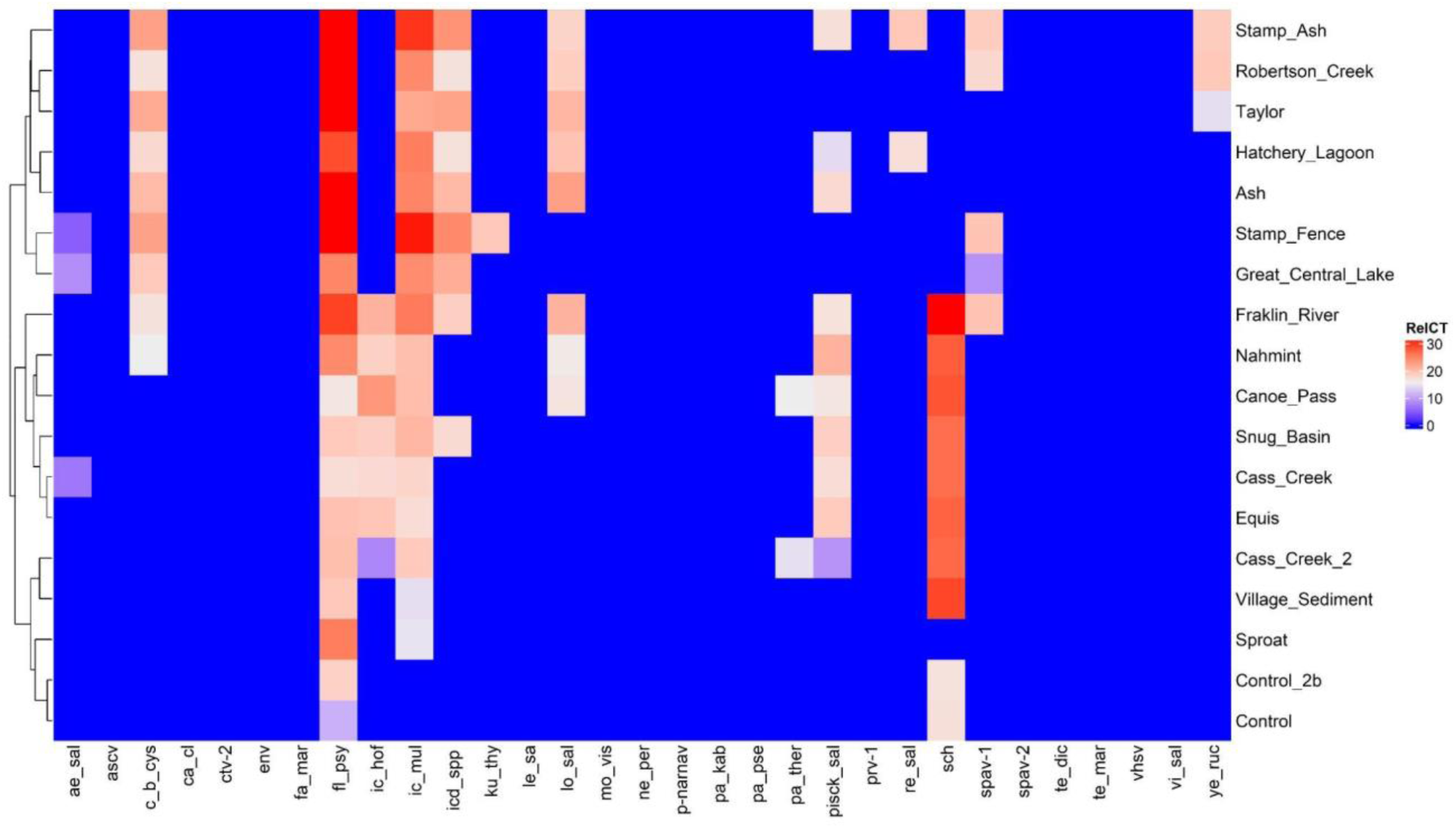
qPCR detections of salmon pathogens, RelCT depicted (45-CT). See Supplementary Table 1 for qPCR assay details.

## Discussion

### Hollow-membrane cartridge filters

HM filters allowed for a substantial increase in filtration volumes compared to equivalent Sterivex filtration cartridges, while mirroring the compact design and contamination protection of Sterivex cartridge filters, enabling efficient handling in the field. However, when comparing the filtration volumes in a near-bottom freshwater system, HM filters were more variable than Sterivex filters, and two HM filters showed highly aberrant filtration volumes. Given that these samples were collected in freshwater, just above the bottom of the river, we cannot dismiss the possibility that the filter that clogged at about 0.5L could have taken up sediment. This specific sampling site was chosen as a challenging environment to reach maximum filtration volumes in a quick and reproducible manner. Freshwater, with high tannins and sediment loading, generally results in lower filtration volumes than in marine environments. However, we suspect that the filter that did not clog until almost 20L may have experienced a disruption of one or more of the 120 HM fibers. In our experience filtering 1000’s of samples with HM filters, the failure rate of filters is generally around ∼2-5%. Speed of filtration for the first 1 L of water was 3-times greater for HM than Sterivex filters across multiple marine environments. When the Sterivex filters were used in diverse marine environments, they showed enhanced variability in filtration speed compared to HM filters, consistent with their lower filtration capacity.

This study did not attempt to provide a head to head comparison of DNA yield recovered from HM and Sterivex filters, e.g. through quantifying nucleic acid recovery from filters by qPCR. Such a quantitative comparison was judged to be not feasible as the samples collected in the field with HM filters (Marine: 5 L; Freshwater 4.50 +/− 0.85L) reached volumes beyond the capacity of sterivex filters. Instead we chose to present the results of the full workup of the HM filters as a case study of the suitability of the HM filters for ecosystem monitoring. Given matching poresize in both types of filters, we assume there to be a quasi-linear relationship between filtration volume and nucleic acid recovery. Accordingly, the higher filtration volume achieved by HM filters should directly translate to increased nucleic acid yields and therefore also boost sensitivity for rare species.

In summary, HM filters are a preferable option for generating large filtration volumes (preferable for metabarcoding and ocean sampling), even in challenging environments. Absolute quantification with qPCR might, similar to established filtration systems, require careful monitoring of filtration volumes and potential adjustment for the latter. Additionally, while cartridge filters can considerably reduce potential for contamination in the field, they require additional steps for nucleic acid extraction, as incomplete nucleic acid recovery from the filtration cartridges of both types might impact absolute quantification.

### Modular eDNA sampling systems

Similar to other systems, like the Smith-Root eDNA sampler, the EZ-eDNA systems allow for compact portable deployment. The EZ-eDNA system is designed around a protective polymer case and uses modular components that can be deployed in a number of different arrangements using different pump and filter configurations. This approach allowed sampling from shore of rivers and lakes, from small vessels in the nearshore environment as well as depth sampling from Niskin bottles in the open ocean. The protected design of the sampling system provides protection from contamination during transport. Additionally, the modular design allows for easy maintenance and repair by the end user. Integrated ozonation further simplified decontamination between samples in the field where recirculation of ozonated water provided generally appropriate decontamination and removed remaining DNA traces from prior samples as assessed with field controls.

### Uu-a-thluk workshop: A case study for salmon management

The eDNA sampling systems described herein were developed first and foremost in order to increase accessibility of this technology for rights and title holders like Indigenous resource management organizations like Uu-a-thluk fisheries. Accordingly, the workshop served as a case study to assess the feasibility of deploying environmental DNA to improve and supplement existing salmon ecosystem management. Within two days of training, all participants of the workshop were proficient in collecting eDNA samples. Further, the eDNA results for salmonid distribution were consistent with conventional assessment methods, while also providing insights into ecosystem composition, distribution of salmon predators and prey as well as salmon pathogens. Additionally, several invasive species could be detected such as Pumpkinseed, Common carp, *Craspedacusta sowerbii*, American bullfrog, Milky Pacific Venus clam (*Compsomyax subdiaphana*), and the Manila clam (*Ruditapes philippinarum*). All these insights were generated within a single day of sampling outperforming conventional surveys in taxonomic depth, breadth and throughput, highlighting the potential of the impending paradigm shift brought about by the environmental DNA revolution. Strategic long term sampling or targeting higher spatial resolution could provide datasets that can be used to investigate the ecological connections that drive salmon ecosystems (e.g. Deeg et al. 2023).

### Aquaculture impacts on eDNA community composition: A cautionary tale

The observation of abundant reads of diverse marine species in freshwater eDNA samples was surprising, but could ultimately be linked to the species contained in the feed used in aquaculture facilities, as detections of these species were highest in samples downstream of the such facilities, and in some cases coincided with the detection of Atlantic salmon that were grown in one of these facilities. Specifically, the sample from Hatchery Lagoon was extraordinarily rich in these reads, making up more than 30% of total reads. This sample was collected right above the sediment at the outflow of Robertson Creek, suggesting that feed associated eDNA is accumulated in the sediment. Similarly, the distribution of salmon pathogens was also impacted by discharge from the aquaculture facilities. However, such detections have to be interpreted with caution as they don’t necessarily represent detections of infective particles. Similar to the detections of marine species, some of these detections could be originating from aquaculture feed. Accordingly, just like the detection of marine fish coming from the feed not suggesting the presence of these fish in freshwater, the detection of said pathogens could represent inactive particles remaining in the aquaculture feed. Nevertheless, the association of pathogen detections with aquaculture discharge in spawning habitat is concerning and warrants further investigation. Together, these observations should serve as a cautionary tale when interpreting community data derived from eDNA that easily can be skewed by industrial activities such as high density animal husbandry.

### Democratization of environmental DNA

Here we describe a novel eDNA sampling system consisting of hollow-membrane cartridge filters and modular water sampling systems. Designed for low complexity and easy adaptation, these systems allow for improved filtration capacities and are suitable for applications employing diverse sampling strategies. With the case study of an eDNA sampling workshop, we demonstrate the adaptation of eDNA sampling by small organizations to inform sustainable salmon management. With eDNA sampling becoming more and more commonplace, the systems provide an accessible option for eDNA sample collection. Looking ahead, improving access to sample processing and analysis will be the next challenges to overcome if eDNA is to become widely adapted by indigenous and coastal communities, and ENGOs.

## Acknowledgements

Uu-a-thluk (https://uuathluk.ca/): Course participants: Shane Sieber, Cody Gus, Erikk Dick-Frank, Robert Watts, Graham Murrell, Christopher Tatoosh, Cameron Tatoosh, Sabrina Crowley, Gemma Macfarlane, Danielle Robertson, Lesley Lauder; Staff: Jim Lane (contact), Alison Wale, Carilia Horning, Sabrina Crowley

Tseshaht Fisheries: Dave Rolston

Pacific Salmon Foundation (PSF): Andrew Bateman

Fisheries and Oceans, Canada (DFO): Lochlan Breckenridge, Rachel Witt, Shannon Adams, Shaye Ryan

Students on Ice (SOI): The course material used in training was developed from funding provided by the SOI Blue Futures program, led by Tara Mascarenhas, to support training in Blue Futures technologies for coastal and Indigenous youth.

## Author contributions

Christoph M. Deeg: Conception, course content, fieldwork, labwork, analysis, writing

Robert G. Saunders: EZ-eDNA System and filter development

Christopher Tam: Fieldwork, labwork (extractions)

Karia Kaukinen: Molecular analysis (qPCR)

Shaorong Li: Molecular analysis (assay design and validation)

Arthur L Bass: Writing

Uu-a-thluk Fisheries: Host workshop and sample collection

Kristina M. Miller: Conception, writing, funding

## Supplementary materials

**Supplementary Table 1:**
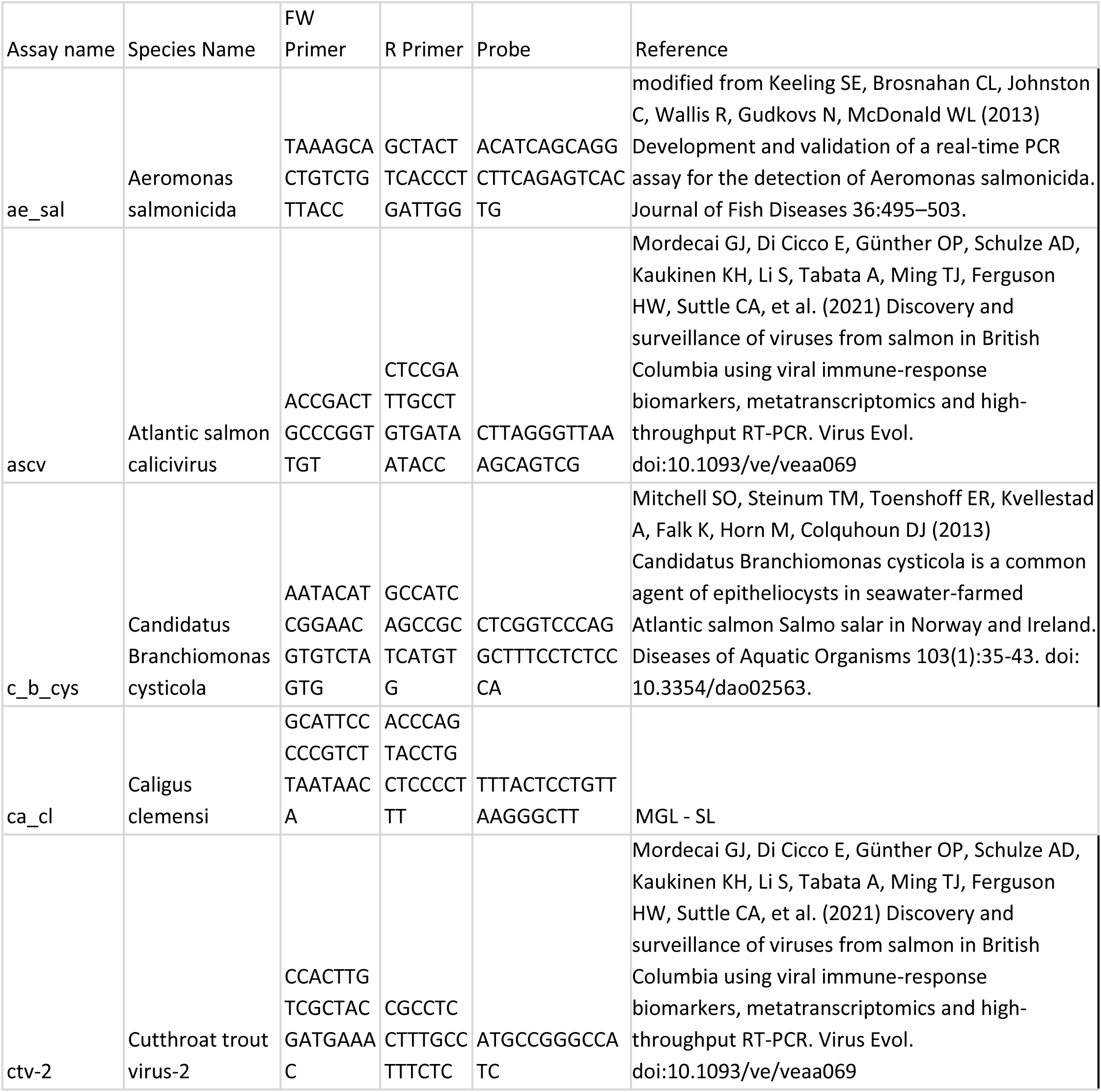
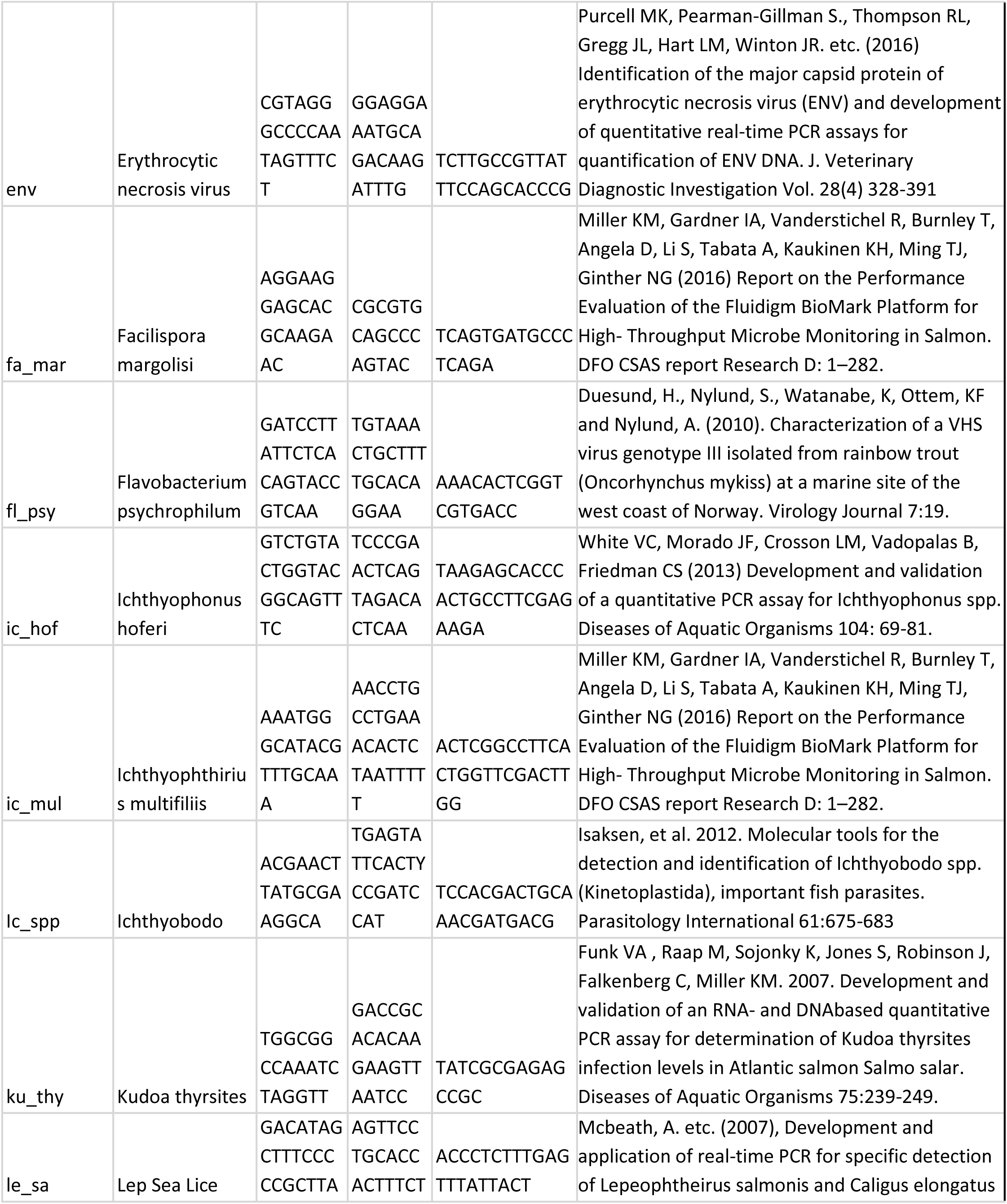
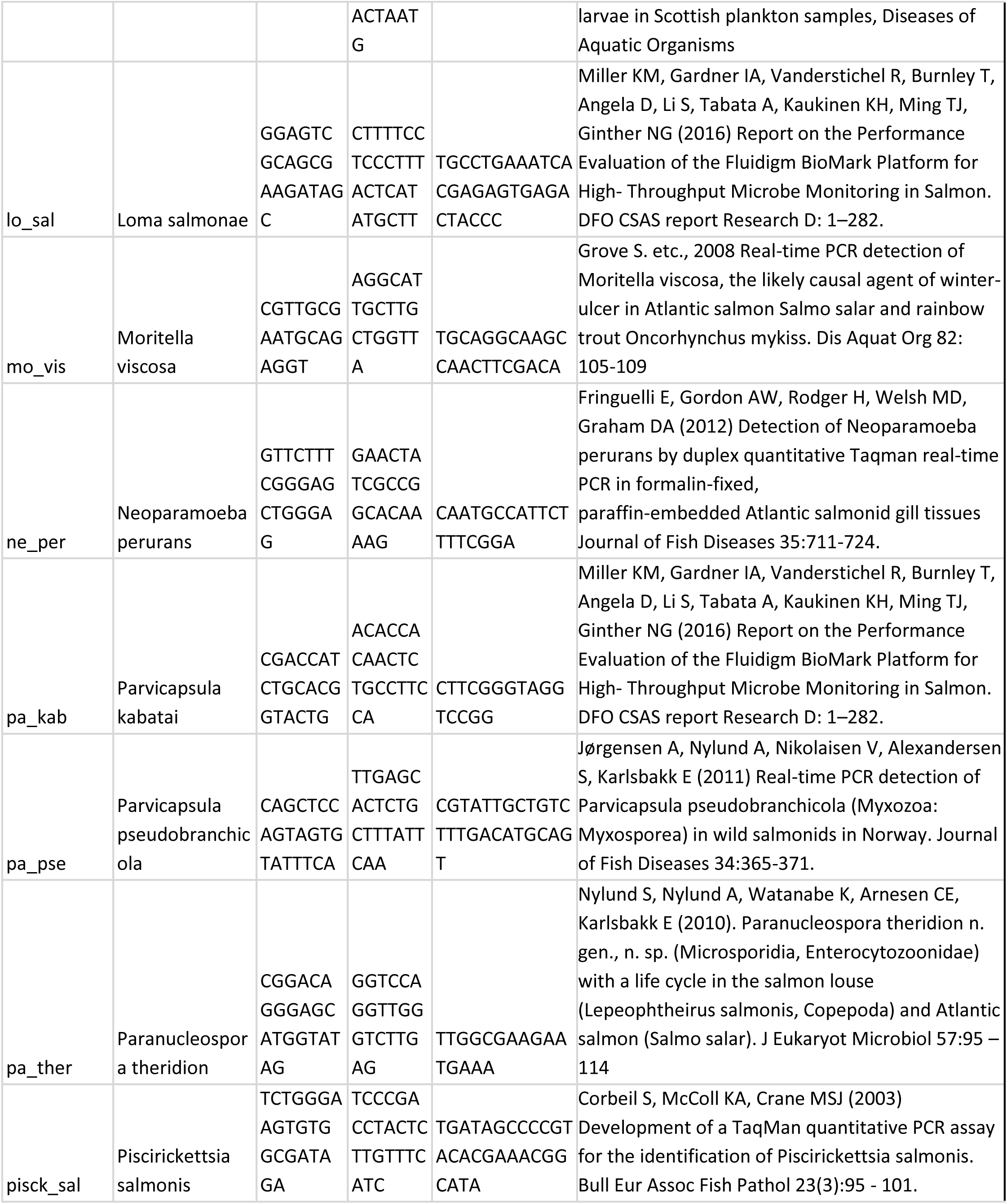
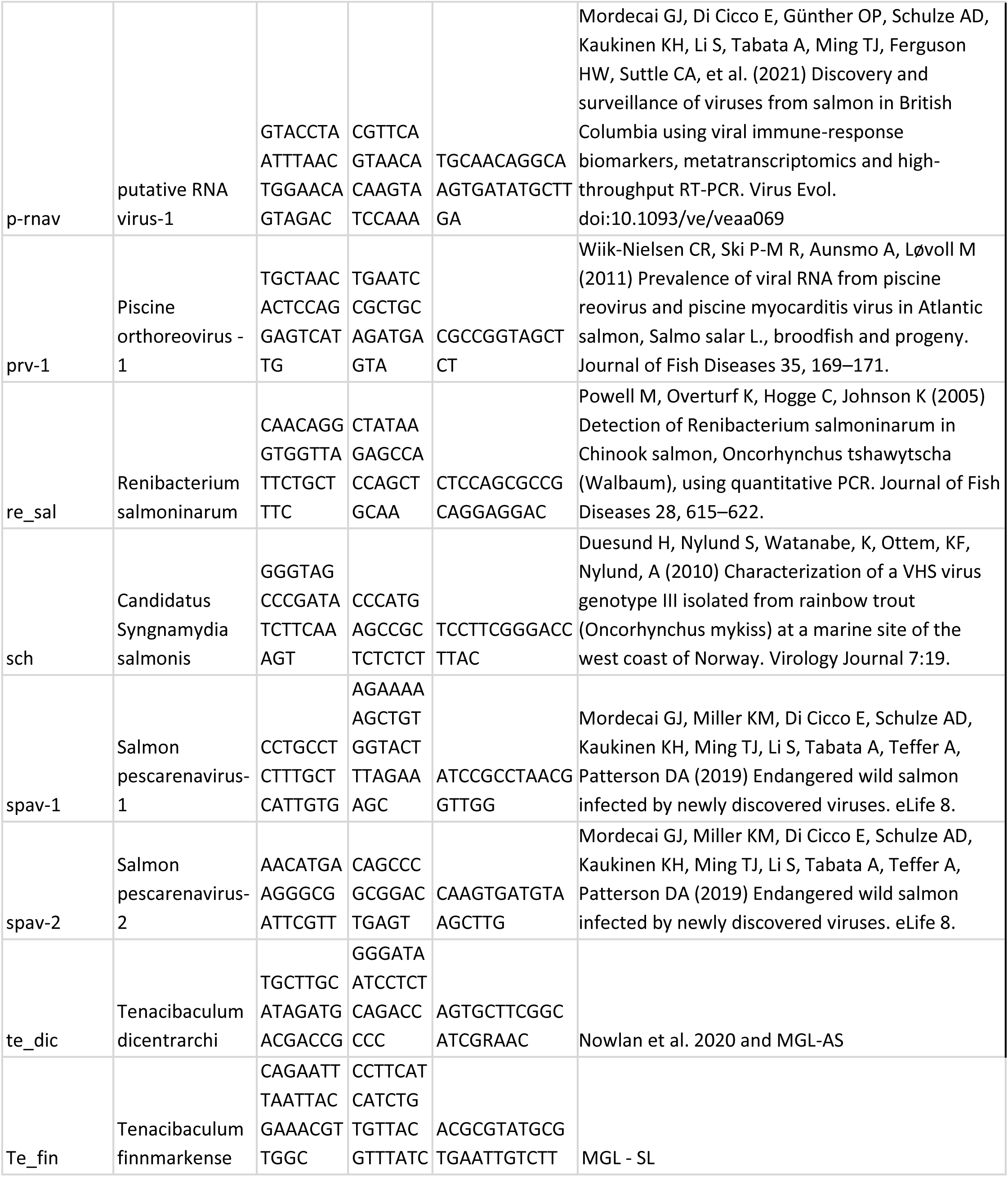
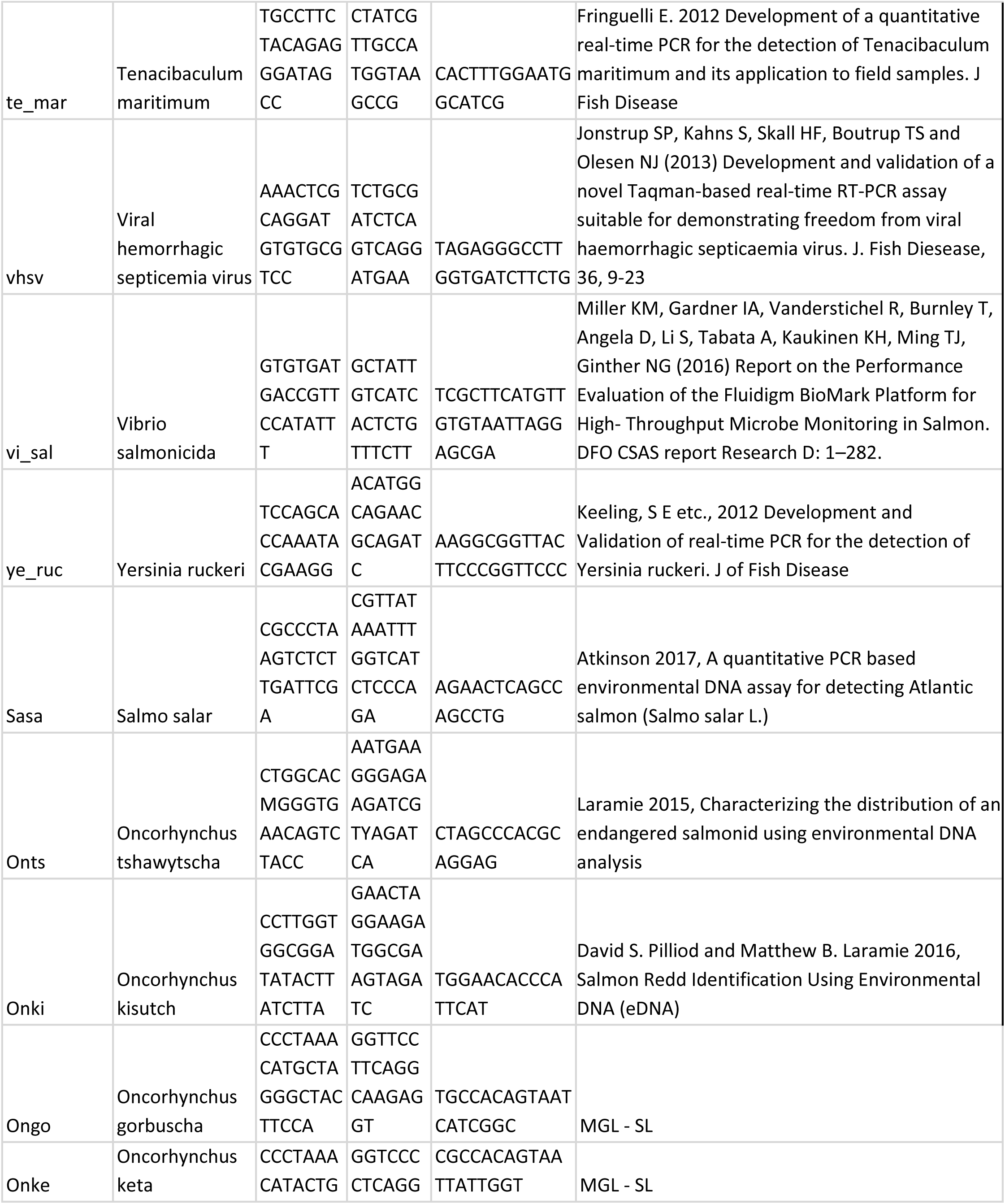
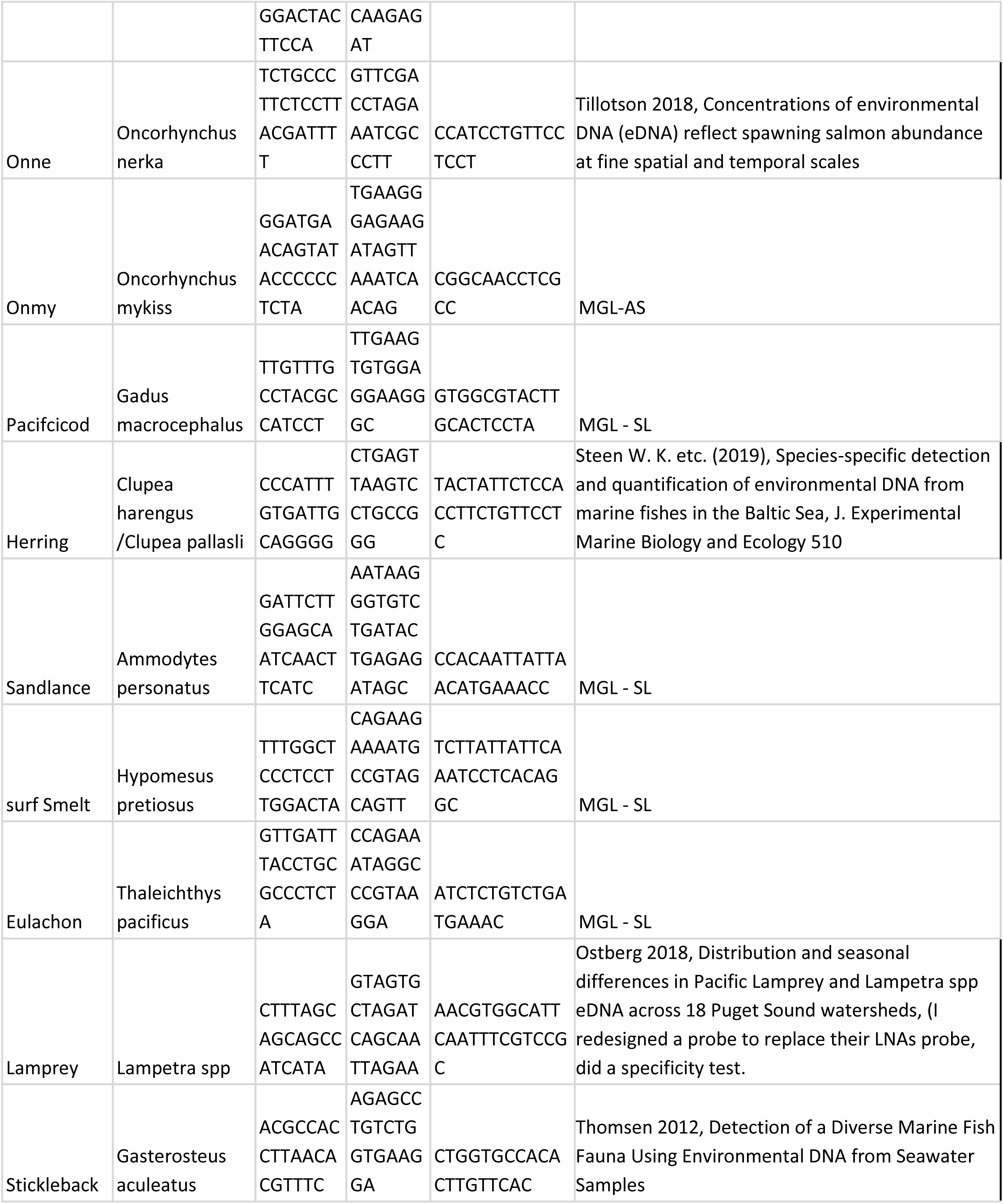
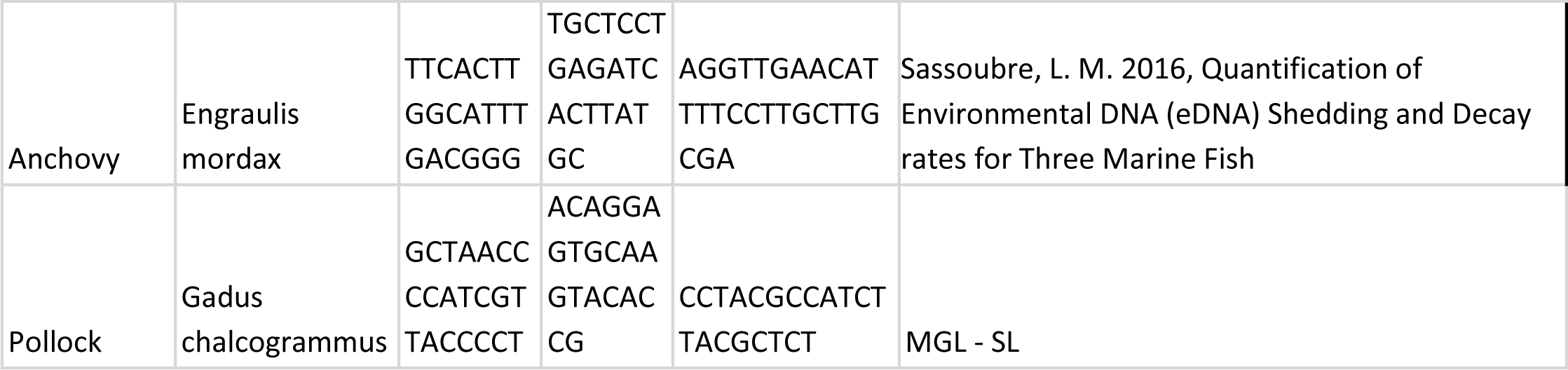
Primer and probes.. Note reference to MGL-SL relates to assays developed in house by our DFO laboratory by technician Shaorong Li (MGL-SL) or Angela Schulze (MGL-AS)

**Supplementary Table 2:**
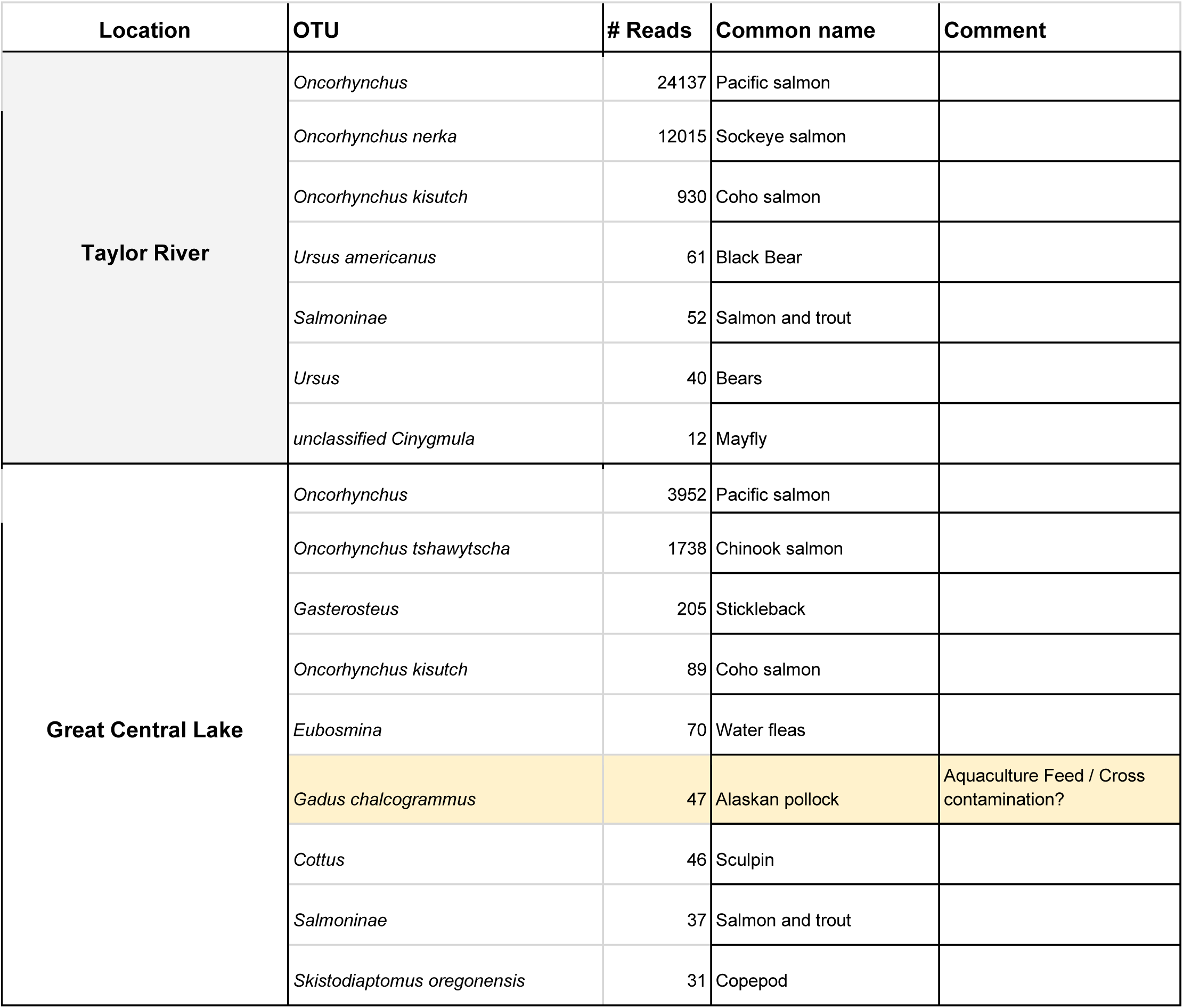
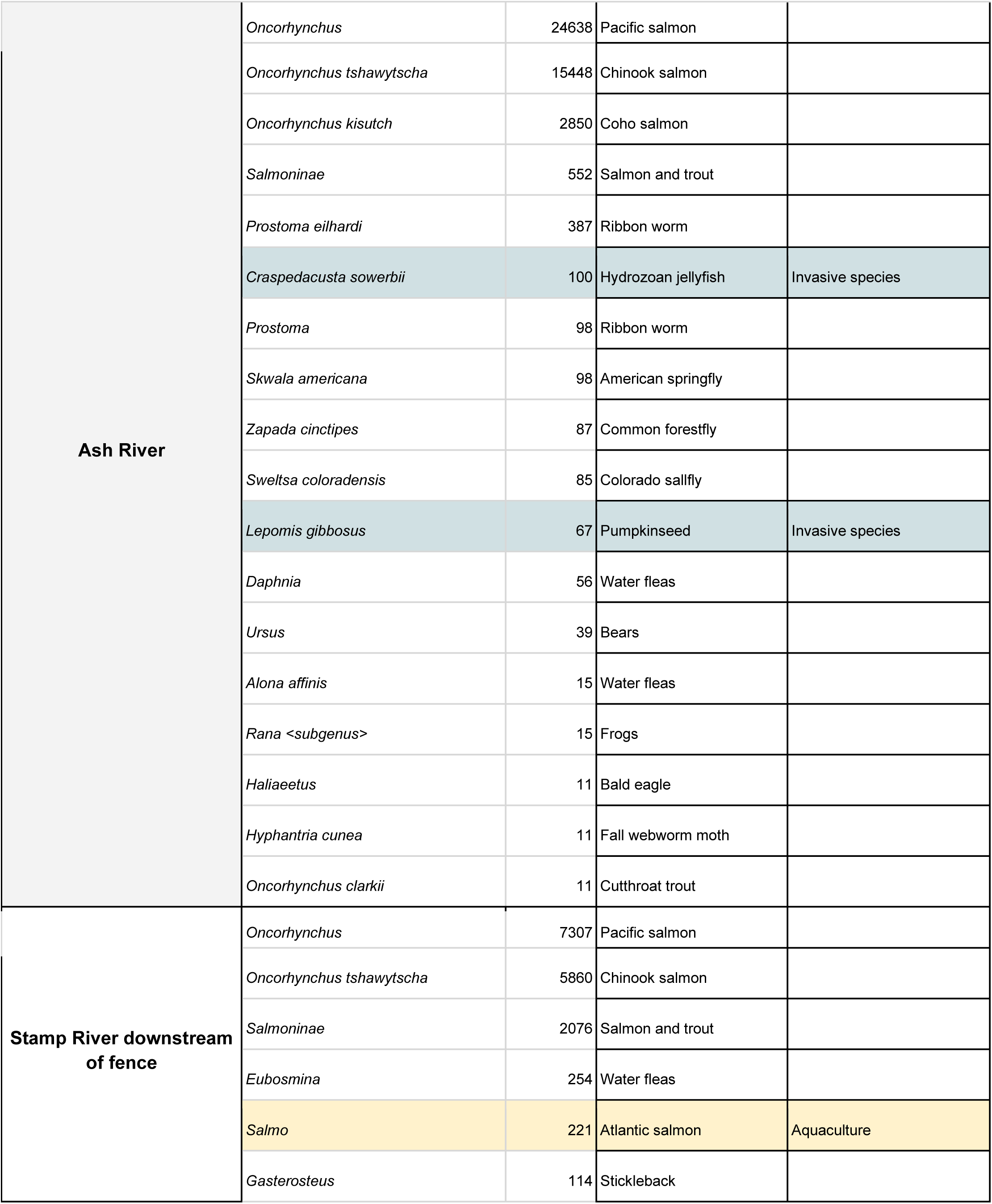
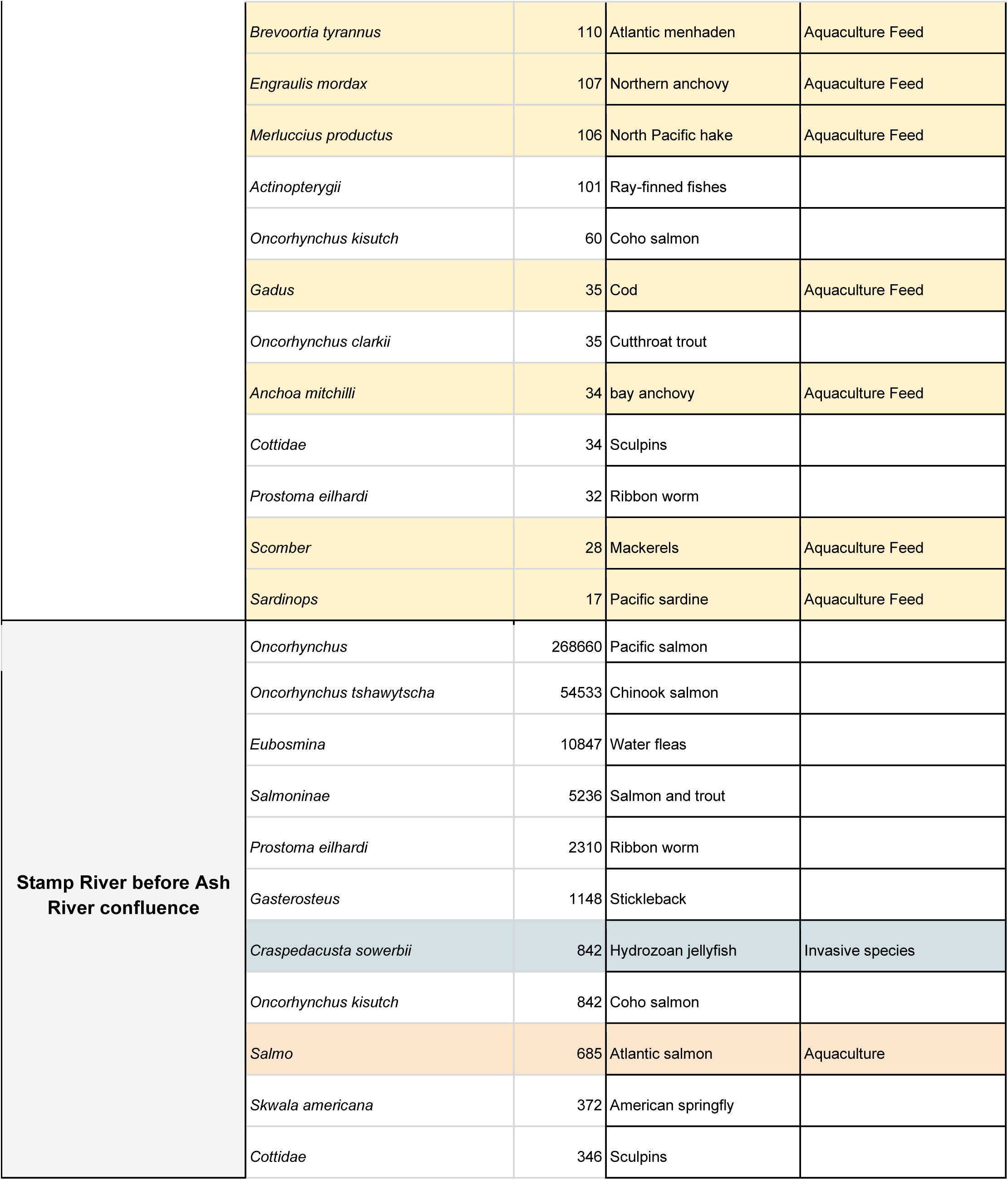
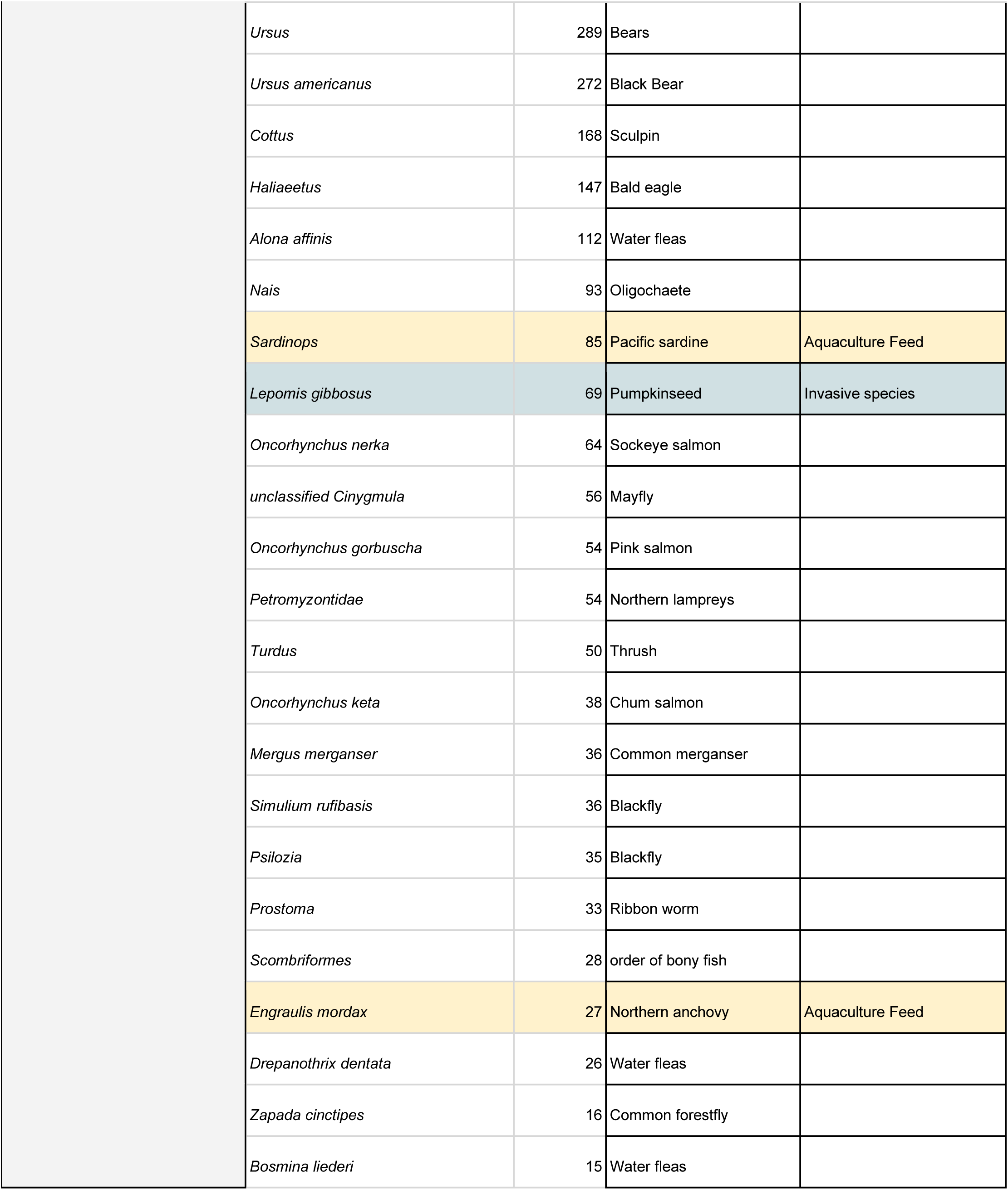
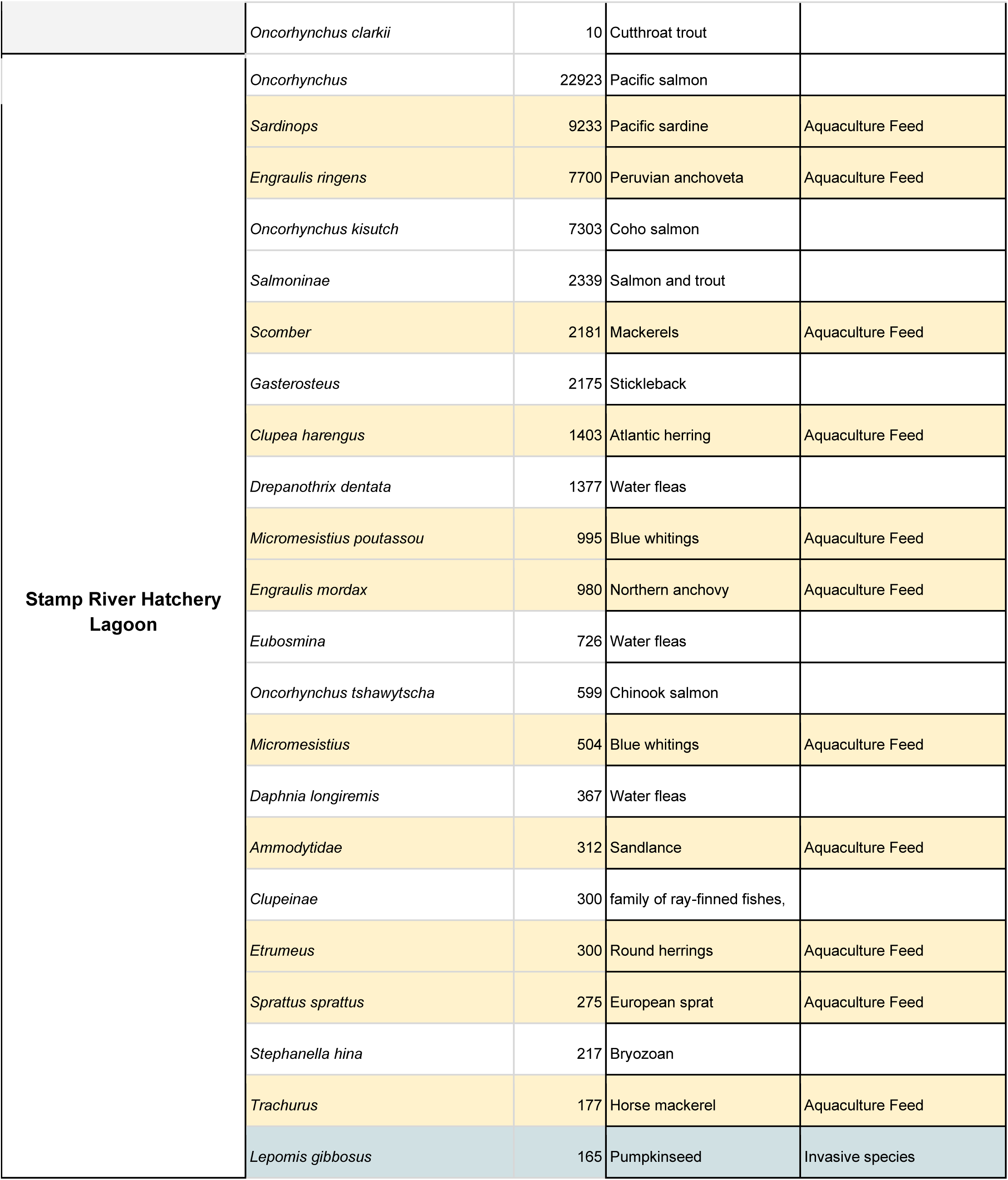
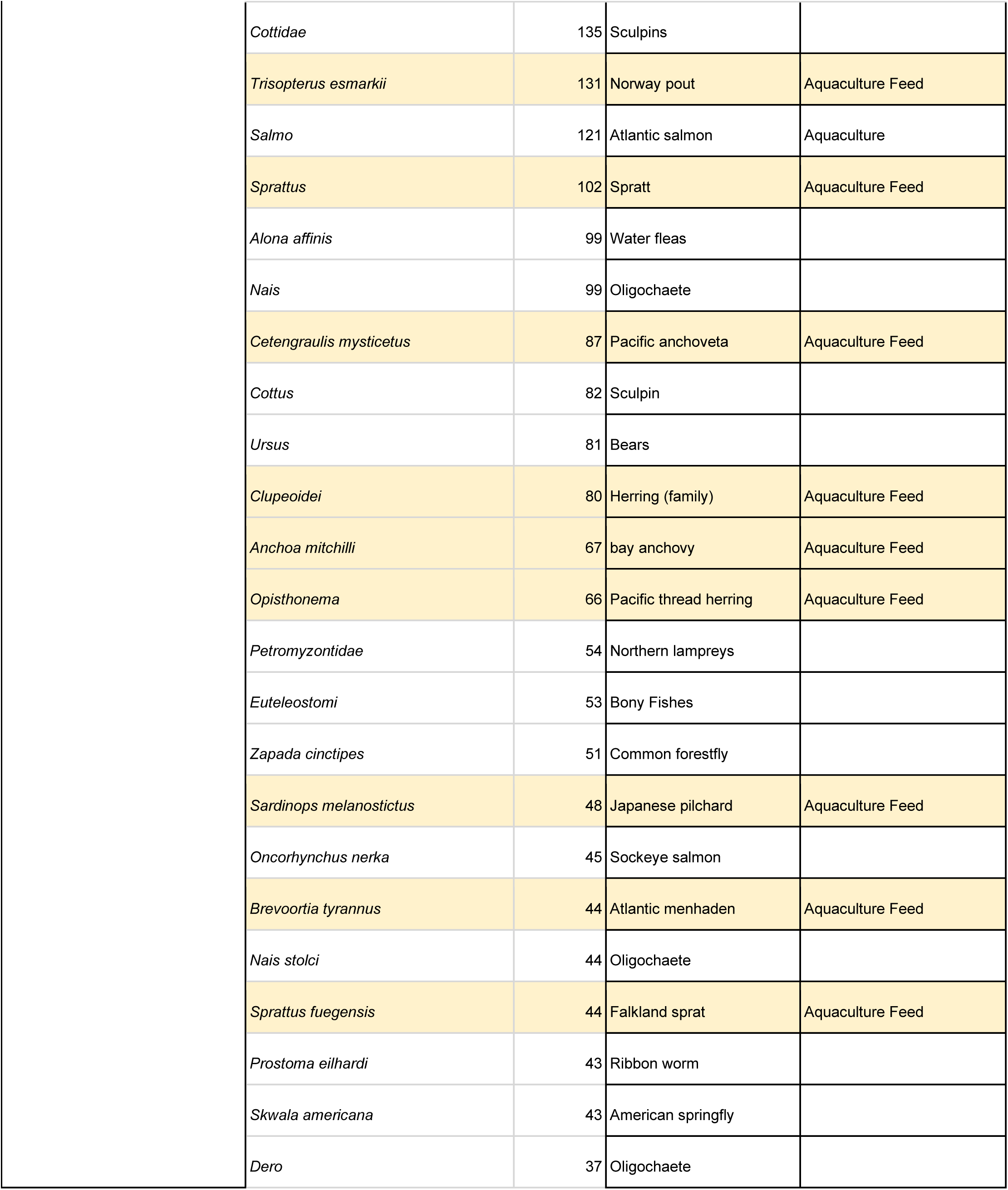
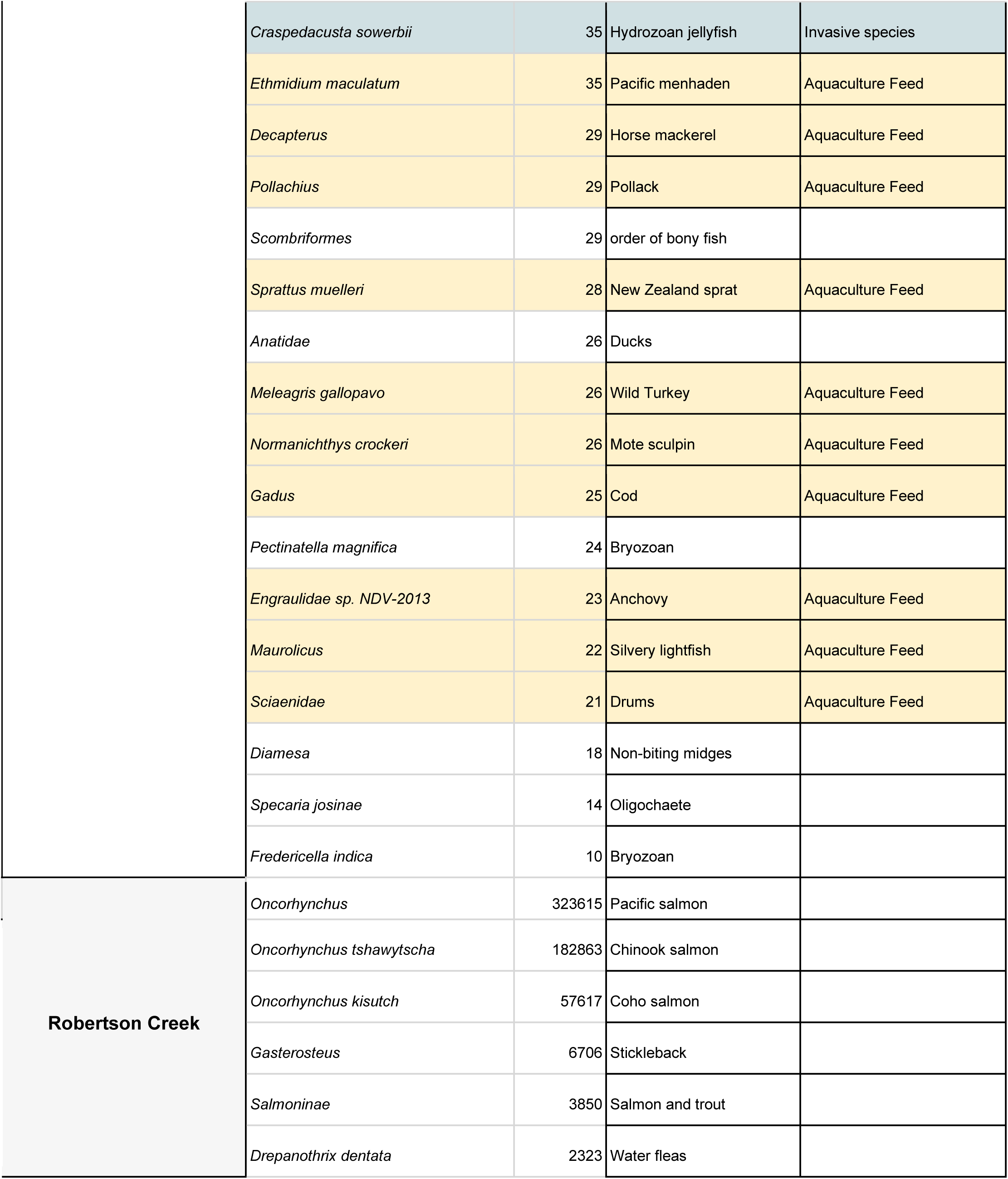
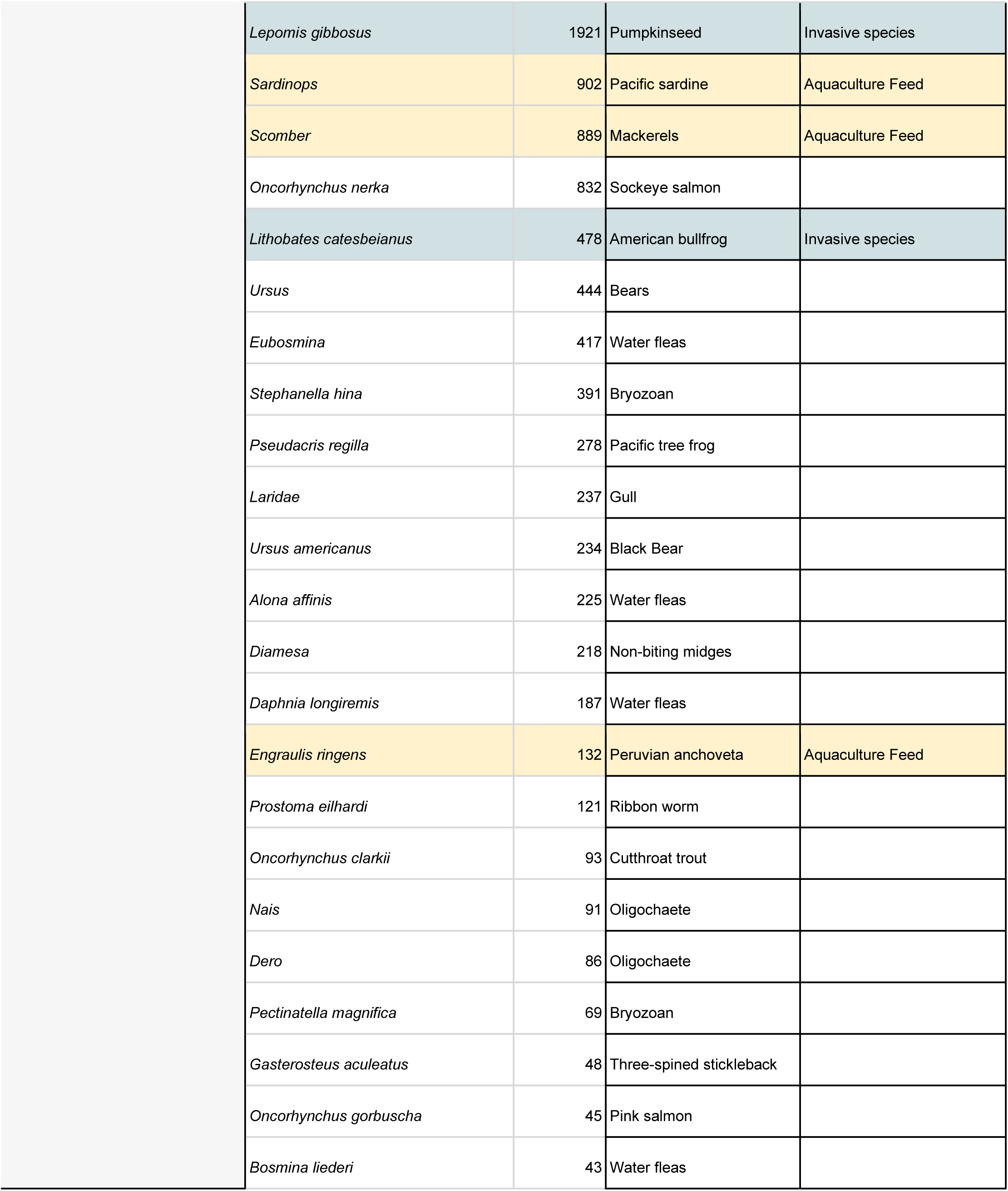
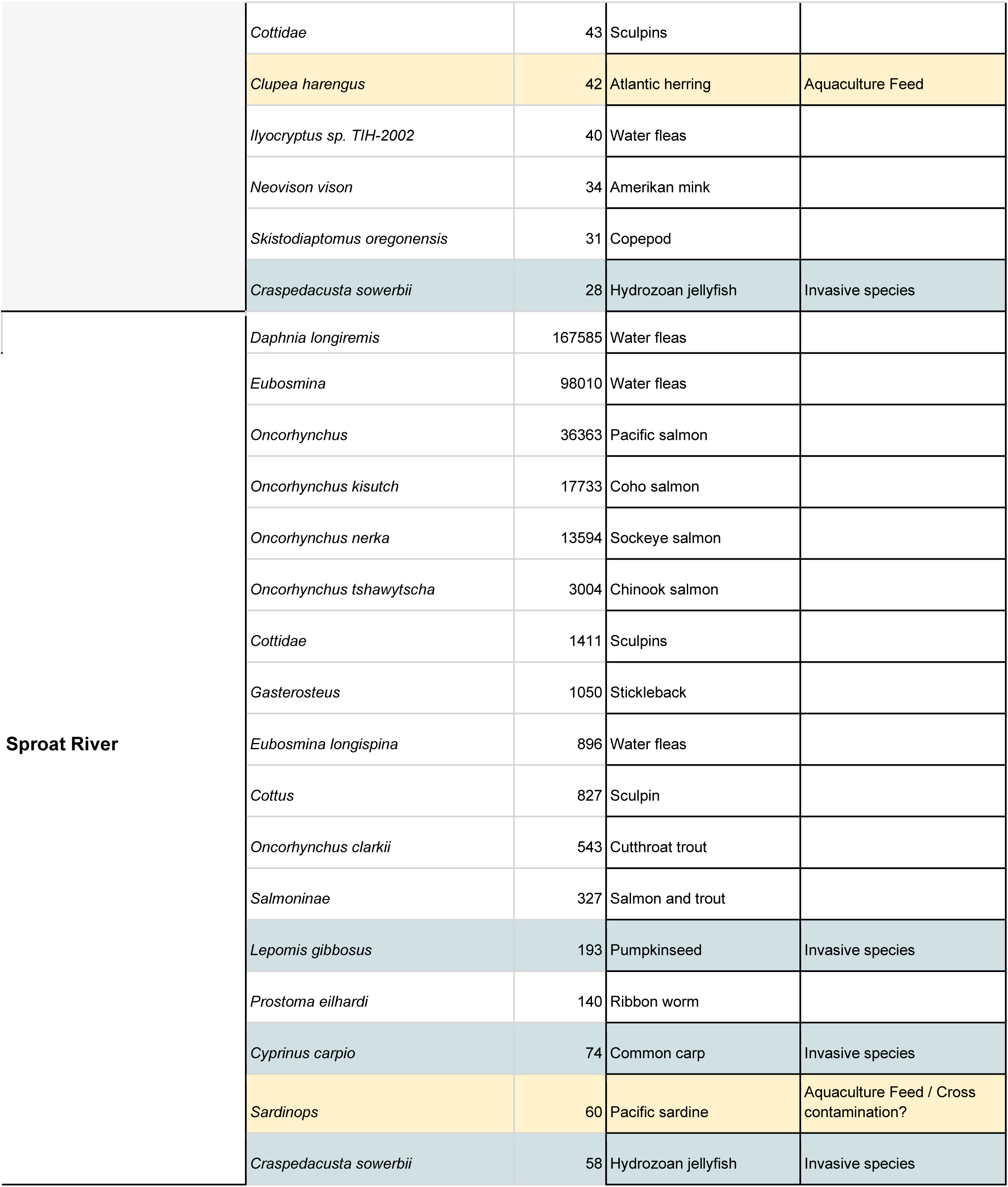
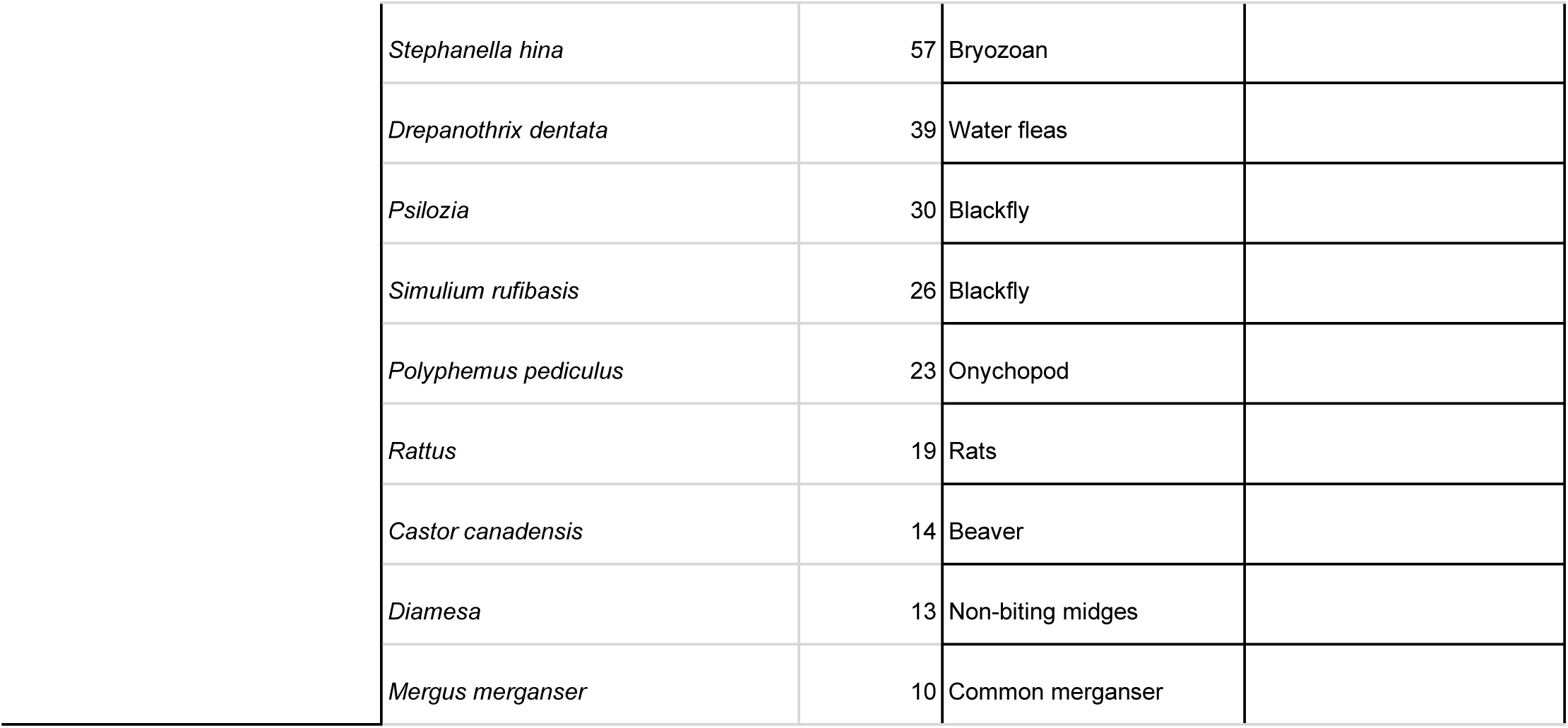
Operational taxonomic unit (OTU) detections in the Somass/Stamp watershed.

| Location | OTU | # Reads | Common name | Comment |
| --- | --- | --- | --- | --- |
| Taylor River | *Oncorhynchus* | 24137 | Pacific salmon |  |
|  | *Oncorhynchus nerka* | 12015 | Sockeye salmon |  |
|  | *Oncorhynchus kisutch* | 930 | Coho salmon |  |
|  | *Ursus americanus* | 61 | Black Bear |  |
|  | *Salmoninae* | 52 | Salmon and trout |  |
|  | *Ursus* | 40 | Bears |  |
|  | *unclassified Cinygmula* | 12 | Mayfly |  |
| Great Central Lake | *Oncorhynchus* | 3952 | Pacific salmon |  |
|  | *Oncorhynchus tshawytscha* | 1738 | Chinook salmon |  |
|  | *Gasterosteus* | 205 | Stickleback |  |
|  | *Oncorhynchus kisutch* | 89 | Coho salmon |  |
|  | *Eubosmina* | 70 | Water fleas |  |
|  | *Gadus chalcogrammus* | 47 | Alaskan pollock | Aquaculture Feed / Cross contamination? |
|  | *Cottus* | 46 | Sculpin |  |
|  | *Salmoninae* | 37 | Salmon and trout |  |
|  | *Skistodiaptomus oregonensis* | 31 | Copepod |  |
| Ash River | *Oncorhynchus* | 24638 | Pacific salmon |  |
|  | *Oncorhynchus tshawytscha* | 15448 | Chinook salmon |  |
|  | *Oncorhynchus kisutch* | 2850 | Coho salmon |  |
|  | *Salmoninae* | 552 | Salmon and trout |  |
|  | *Prostoma eilhardi* | 387 | Ribbon worm |  |
|  | *Craspedacusta sowerbii* | 100 | Hydrozoan jellyfish | Invasive species |
|  | *Prostoma* | 98 | Ribbon worm |  |
|  | *Skwala americana* | 98 | American springfly |  |
|  | *Zapada cinctipes* | 87 | Common forestfly |  |
|  | *Sweltsa coloradensis* | 85 | Colorado sallfly |  |
|  | *Lepomis gibbosus* | 67 | Pumpkinseed | Invasive species |
|  | *Daphnia* | 56 | Water fleas |  |
|  | *Ursus* | 39 | Bears |  |
|  | *Alona affinis* | 15 | Water fleas |  |
|  | *Rana <subgenus>* | 15 | Frogs |  |
|  | *Haliaeetus* | 11 | Bald eagle |  |
|  | *Hyphantria cunea* | 11 | Fall webworm moth |  |
|  | *Oncorhynchus clarkii* | 11 | Cutthroat trout |  |
| Stamp River downstream of fence | *Oncorhynchus* | 7307 | Pacific salmon |  |
|  | *Oncorhynchus tshawytscha* | 5860 | Chinook salmon |  |
|  | *Salmoninae* | 2076 | Salmon and trout |  |
|  | *Eubosmina* | 254 | Water fleas |  |
|  | *Salmo* | 221 | Atlantic salmon | Aquaculture |
|  | *Gasterosteus* | 114 | Stickleback |  |
|  | *Brevoortia tyrannus* | 110 | Atlantic menhaden | Aquaculture Feed |
|  | *Engraulis mordax* | 107 | Northern anchovy | Aquaculture Feed |
|  | *Merluccius productus* | 106 | North Pacific hake | Aquaculture Feed |
|  | *Actinopterygii* | 101 | Ray-finned fishes |  |
|  | *Oncorhynchus kisutch* | 60 | Coho salmon |  |
|  | *Gadus* | 35 | Cod | Aquaculture Feed |
|  | *Oncorhynchus clarkii* | 35 | Cutthroat trout |  |
|  | *Anchoa mitchilli* | 34 | bay anchovy | Aquaculture Feed |
|  | *Cottidae* | 34 | Sculpins |  |
|  | *Prostoma eilhardi* | 32 | Ribbon worm |  |
|  | *Scomber* | 28 | Mackerels | Aquaculture Feed |
|  | *Sardinops* | 17 | Pacific sardine | Aquaculture Feed |
| Stamp River before Ash<br>River confluence | *Oncorhynchus* | 268660 | Pacific salmon |  |
|  | *Oncorhynchus tshawytscha* | 54533 | Chinook salmon |  |
|  | *Eubosmina* | 10847 | Water fleas |  |
|  | *Salmoninae* | 5236 | Salmon and trout |  |
|  | *Prostoma eilhardi* | 2310 | Ribbon worm |  |
|  | *Gasterosteus* | 1148 | Stickleback |  |
|  | *Craspedacusta sowerbii* | 842 | Hydrozoan jellyfish | Invasive species |
|  | *Oncorhynchus kisutch* | 842 | Coho salmon |  |
|  | *Salmo* | 685 | Atlantic salmon | Aquaculture |
|  | *Skwala americana* | 372 | American springfly |  |
|  | *Cottidae* | 346 | Sculpins |  |
|  | *Ursus* | 289 | Bears |  |
|  | *Ursus americanus* | 272 | Black Bear |  |
|  | *Cottus* | 168 | Sculpin |  |
|  | *Haliaeetus* | 147 | Bald eagle |  |
|  | *Alona affinis* | 112 | Water fleas |  |
|  | *Nais* | 93 | Oligochaete |  |
|  | *Sardinops* | 85 | Pacific sardine | Aquaculture Feed |
|  | *Lepomis gibbosus* | 69 | Pumpkinseed | Invasive species |
|  | *Oncorhynchus nerka* | 64 | Sockeye salmon |  |
|  | *unclassified Cinygmula* | 56 | Mayfly |  |
|  | *Oncorhynchus gorbuscha* | 54 | Pink salmon |  |
|  | *Petromyzontidae* | 54 | Northern lampreys |  |
|  | *Turdus* | 50 | Thrush |  |
|  | *Oncorhynchus keta* | 38 | Chum salmon |  |
|  | *Mergus merganser* | 36 | Common merganser |  |
|  | *Simulium rufibasis* | 36 | Blackfly |  |
|  | *Psilozia* | 35 | Blackfly |  |
|  | *Prostoma* | 33 | Ribbon worm |  |
|  | *Scombriformes* | 28 | order of bony fish |  |
|  | *Engraulis mordax* | 27 | Northern anchovy | Aquaculture Feed |
|  | *Drepanothrix dentata* | 26 | Water fleas |  |
|  | *Zapada cinctipes* | 16 | Common forestfly |  |
|  | *Bosmina luederi* | 15 | Water fleas |  |
|  | *Oncorhynchus clarkii* | 10 | Cutthroat trout |  |
| Stamp River Hatchery<br>Lagoon | *Oncorhynchus* | 22923 | Pacific salmon |  |
|  | *Sardinops* | 9233 | Pacific sardine | Aquaculture Feed |
|  | *Engraulis ringens* | 7700 | Peruvian anchoveta | Aquaculture Feed |
|  | *Oncorhynchus kisutch* | 7303 | Coho salmon |  |
|  | *Salmoninae* | 2339 | Salmon and trout |  |
|  | *Scomber* | 2181 | Mackerels | Aquaculture Feed |
|  | *Gasterosteus* | 2175 | Stickleback |  |
|  | *Clupea harengus* | 1403 | Atlantic herring | Aquaculture Feed |
|  | *Drepanothrix dentata* | 1377 | Water fleas |  |
|  | *Micromesistius poutassou* | 995 | Blue whittings | Aquaculture Feed |
|  | *Engraulis mordax* | 980 | Northern anchovy | Aquaculture Feed |
|  | *Eubosmina* | 726 | Water fleas |  |
|  | *Oncorhynchus tshawytscha* | 599 | Chinook salmon |  |
|  | *Micromesistius* | 504 | Blue whittings | Aquaculture Feed |
|  | *Daphnia longiremis* | 367 | Water fleas |  |
|  | *Ammodytidae* | 312 | Sandlance | Aquaculture Feed |
|  | *Clupeinae* | 300 | family of ray-finned fishes, |  |
|  | *Etrumeus* | 300 | Round herrings | Aquaculture Feed |
|  | *Sprattus sprattus* | 275 | European sprat | Aquaculture Feed |
|  | *Stephanella hina* | 217 | Bryozoan |  |
|  | *Trachurus* | 177 | Horse mackerel | Aquaculture Feed |
|  | *Lepomis gibbosus* | 165 | Pumpkinseed | Invasive species |

|  |  |  |  |
| --- | --- | --- | --- |
| *Cottidae* | 135 | Sculpins |  |
| *Trisopterus esmarkii* | 131 | Norway pout | Aquaculture Feed |
| *Salmo* | 121 | Atlantic salmon | Aquaculture |
| *Sprattus* | 102 | Spratt | Aquaculture Feed |
| *Alona affinis* | 99 | Water fleas |  |
| *Nais* | 99 | Oligochaete |  |
| *Cetengraulis mysticetus* | 87 | Pacific anchoveta | Aquaculture Feed |
| *Cottus* | 82 | Sculpin |  |
| *Ursus* | 81 | Bears |  |
| *Clupeoidei* | 80 | Herring (family) | Aquaculture Feed |
| *Anchoa mitchilli* | 67 | bay anchovy | Aquaculture Feed |
| *Opisthonema* | 66 | Pacific thread herring | Aquaculture Feed |
| *Petromyzontidae* | 54 | Northern lampreys |  |
| *Euteleostomi* | 53 | Bony Fishes |  |
| *Zapada cinctipes* | 51 | Common forestfly |  |
| *Sardinops melanostictus* | 48 | Japanese pilchard | Aquaculture Feed |
| *Oncorhynchus nerka* | 45 | Sockeye salmon |  |
| *Brevoortia tyrannus* | 44 | Atlantic menhaden | Aquaculture Feed |
| *Nais stolci* | 44 | Oligochaete |  |
| *Sprattus fuegensis* | 44 | Falkland sprat | Aquaculture Feed |
| *Prostoma eilhardi* | 43 | Ribbon worm |  |
| *Skwala americana* | 43 | American springfly |  |
| *Dero* | 37 | Oligochaete |  |
|  | *Craspedacusta sowerbii* | 35 | Hydrozoan jellyfish | Invasive species |
|  | *Ethmidium maculatum* | 35 | Pacific menhaden | Aquaculture Feed |
|  | *Decapterus* | 29 | Horse mackerel | Aquaculture Feed |
|  | *Pollachius* | 29 | Pollack | Aquaculture Feed |
|  | *Scombriformes* | 29 | order of bony fish |  |
|  | *Sprattus muelleri* | 28 | New Zealand sprat | Aquaculture Feed |
|  | *Anatidae* | 26 | Ducks |  |
|  | *Meleagris gallopavo* | 26 | Wild Turkey | Aquaculture Feed |
|  | *Normanichthys crockeri* | 26 | Mote sculpin | Aquaculture Feed |
|  | *Gadus* | 25 | Cod | Aquaculture Feed |
|  | *Pectinatella magnifica* | 24 | Bryozoan |  |
|  | *Engraulidae sp. NDV-2013* | 23 | Anchovy | Aquaculture Feed |
|  | *Maurollicus* | 22 | Silvery lightfish | Aquaculture Feed |
|  | *Sciaenidae* | 21 | Drums | Aquaculture Feed |
|  | *Diamesa* | 18 | Non-biting midges |  |
|  | *Specaria josinae* | 14 | Oligochaete |  |
|  | *Fredericella indica* | 10 | Bryozoan |  |
| Robertson Creek | *Oncorhynchus* | 323615 | Pacific salmon |  |
|  | *Oncorhynchus tshawytscha* | 182863 | Chinook salmon |  |
|  | *Oncorhynchus kisutch* | 57617 | Coho salmon |  |
|  | *Gasterosteus* | 6706 | Stickleback |  |
|  | *Salmoninae* | 3850 | Salmon and trout |  |
|  | *Drepanothrix dentata* | 2323 | Water fleas |  |

|  |  |  |  |
| --- | --- | --- | --- |
| *Lepomis gibbosus* | 1921 | Pumpkinseed | Invasive species |
| *Sardinops* | 902 | Pacific sardine | Aquaculture Feed |
| *Scomber* | 889 | Mackerels | Aquaculture Feed |
| *Oncorhynchus nerka* | 832 | Sockeye salmon |  |
| *Lithobates catesbeianus* | 478 | American bullfrog | Invasive species |
| *Ursus* | 444 | Bears |  |
| *Eubosmina* | 417 | Water fleas |  |
| *Stephanella hina* | 391 | Bryozoan |  |
| *Pseudacris regilla* | 278 | Pacific tree frog |  |
| *Laridae* | 237 | Gull |  |
| *Ursus americanus* | 234 | Black Bear |  |
| *Alona affinis* | 225 | Water fleas |  |
| *Diamesa* | 218 | Non-biting midges |  |
| *Daphnia longiremis* | 187 | Water fleas |  |
| *Engraulis ringens* | 132 | Peruvian anchoveta | Aquaculture Feed |
| *Prostoma eilhardi* | 121 | Ribbon worm |  |
| *Oncorhynchus clarkii* | 93 | Cutthroat trout |  |
| *Nais* | 91 | Oligochaete |  |
| *Dero* | 86 | Oligochaete |  |
| *Pectinatella magnifica* | 69 | Bryozoan |  |
| *Gasterosteus aculeatus* | 48 | Three-spined stickleback |  |
| *Oncorhynchus gorbuscha* | 45 | Pink salmon |  |
| *Bosmina luederi* | 43 | Water fleas |  |
|  | *Cottidae* | 43 | Sculpins |  |
|  | *Clupea harengus* | 42 | Atlantic herring | Aquaculture Feed |
|  | *Ilyocryptus* sp. TIH-2002 | 40 | Water fleas |  |
|  | *Neovison vison* | 34 | Amerikan mink |  |
|  | *Skistodiaptomus oregonensis* | 31 | Copepod |  |
|  | *Craspedacusta sowerbii* | 28 | Hydrozoan jellyfish | Invasive species |
| Sproat River | *Daphnia longiremis* | 167585 | Water fleas |  |
|  | *Eubosmina* | 98010 | Water fleas |  |
|  | *Oncorhynchus* | 36363 | Pacific salmon |  |
|  | *Oncorhynchus kisutch* | 17733 | Coho salmon |  |
|  | *Oncorhynchus nerka* | 13594 | Sockeye salmon |  |
|  | *Oncorhynchus tshawytscha* | 3004 | Chinook salmon |  |
|  | *Cottidae* | 1411 | Sculpins |  |
|  | *Gasterosteus* | 1050 | Stickleback |  |
|  | *Eubosmina longispina* | 896 | Water fleas |  |
|  | *Cottus* | 827 | Sculpin |  |
|  | *Oncorhynchus clarkii* | 543 | Cutthroat trout |  |
|  | *Salmoninae* | 327 | Salmon and trout |  |
|  | *Lepomis gibbosus* | 193 | Pumpkinseed | Invasive species |
|  | *Prostoma eilhardi* | 140 | Ribbon worm |  |
|  | *Cyprinus carpio* | 74 | Common carp | Invasive species |
|  | *Sardinops* | 60 | Pacific sardine | Aquaculture Feed / Cross contamination? |
|  | *Craspedacusta sowerbii* | 58 | Hydrozoan jellyfish | Invasive species |

|  |  |  |
| --- | --- | --- |
| *Stephanella hina* | 57 | Bryozoan |
| *Drepanothrix dentata* | 39 | Water fleas |
| *Psilozia* | 30 | Blackfly |
| *Simulium rufibasis* | 26 | Blackfly |
| *Polyphemus pediculus* | 23 | Onychopod |
| *Rattus* | 19 | Rats |
| *Castor canadensis* | 14 | Beaver |
| *Diamesa* | 13 | Non-biting midges |
| *Mergus merganser* | 10 | Common merganser |

**Supplementary Table 3:**
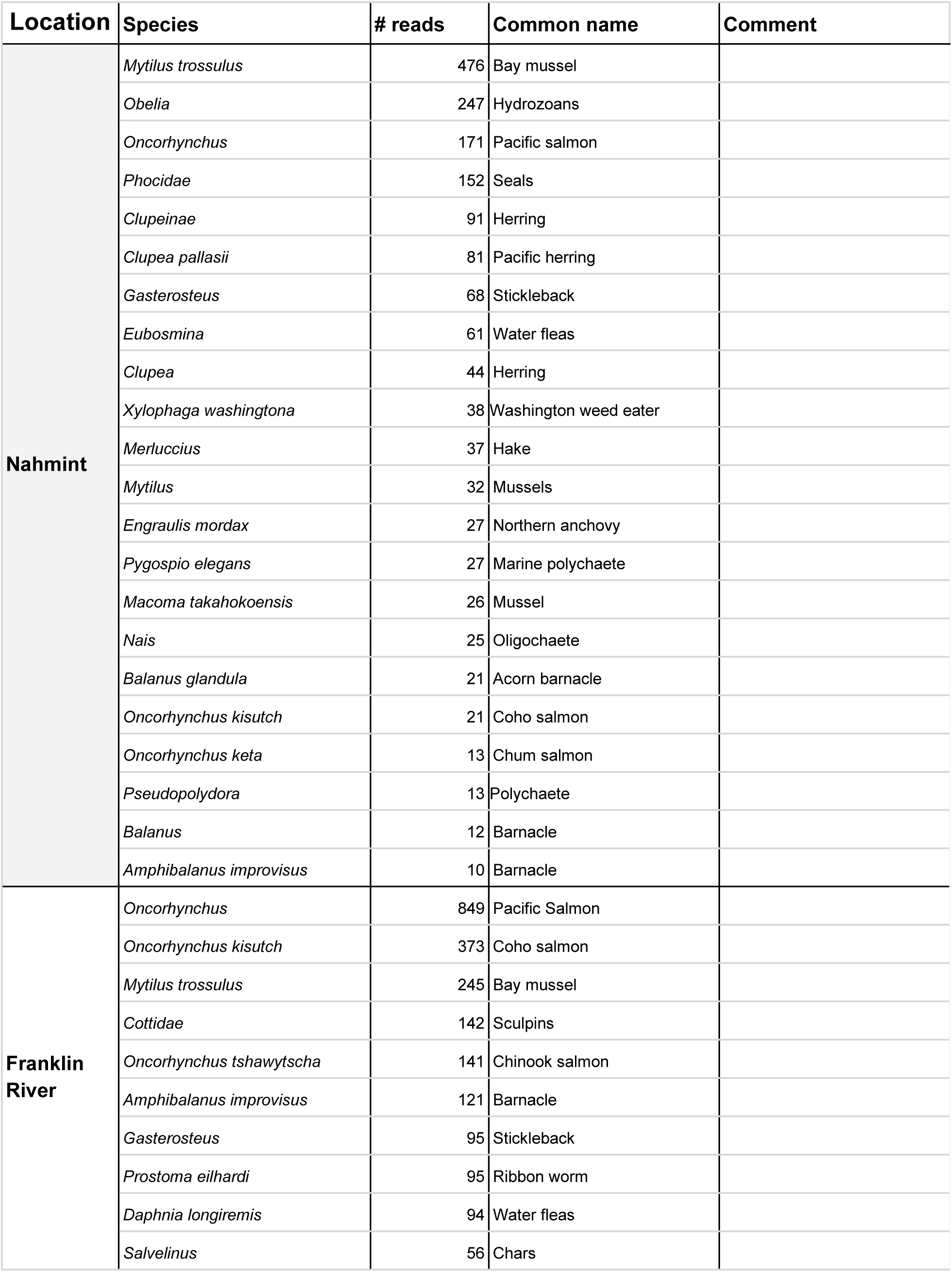
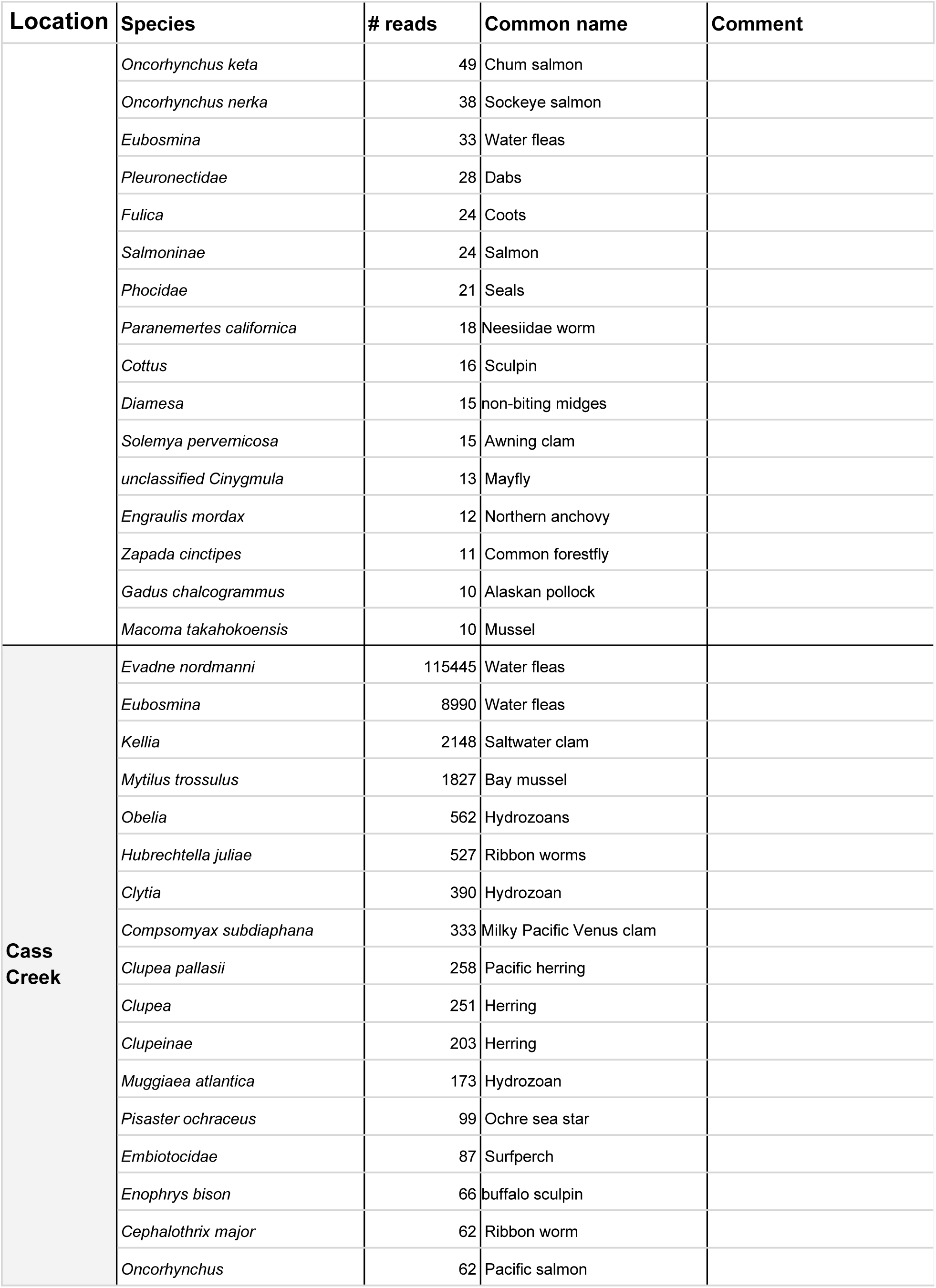
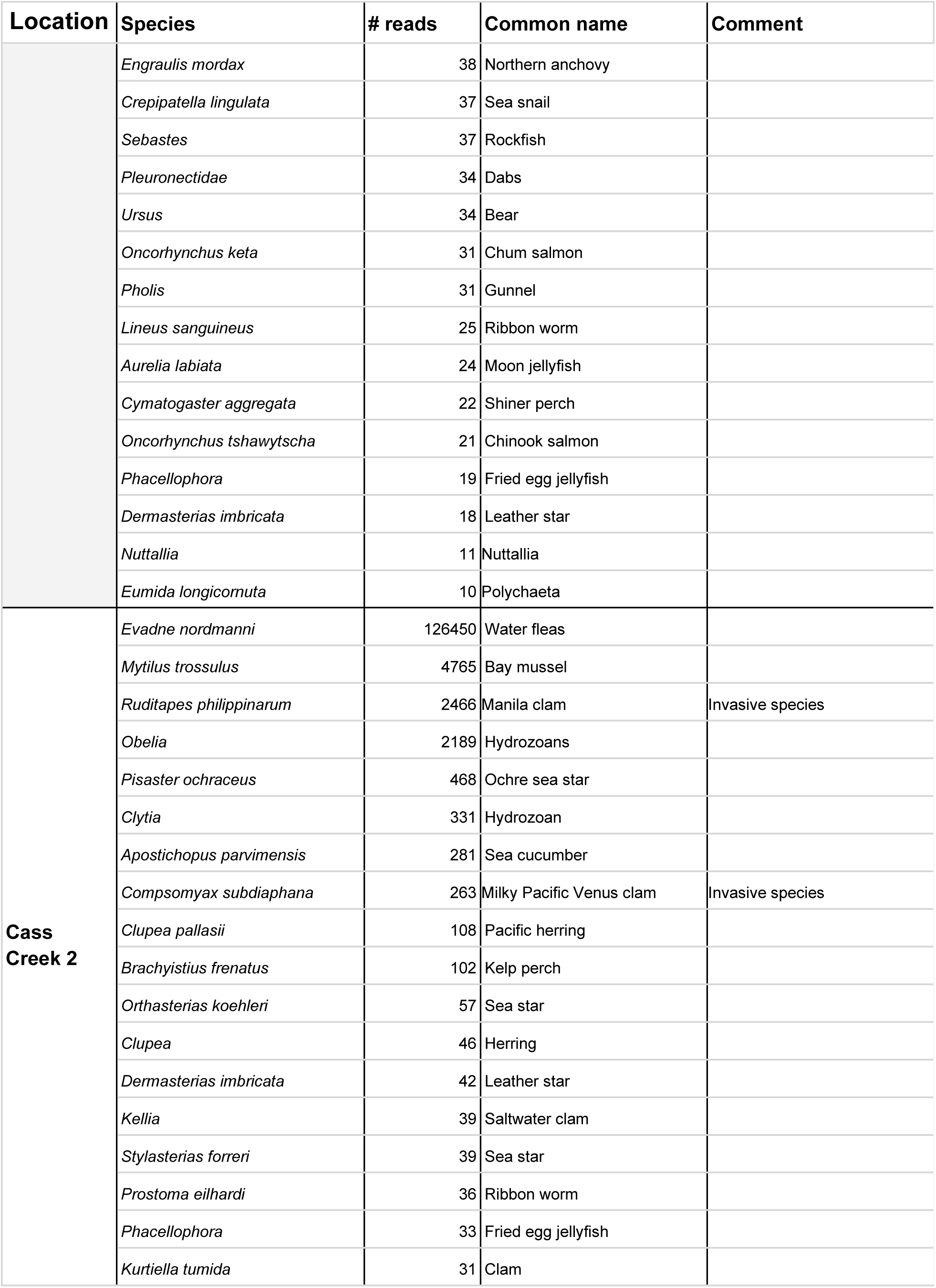
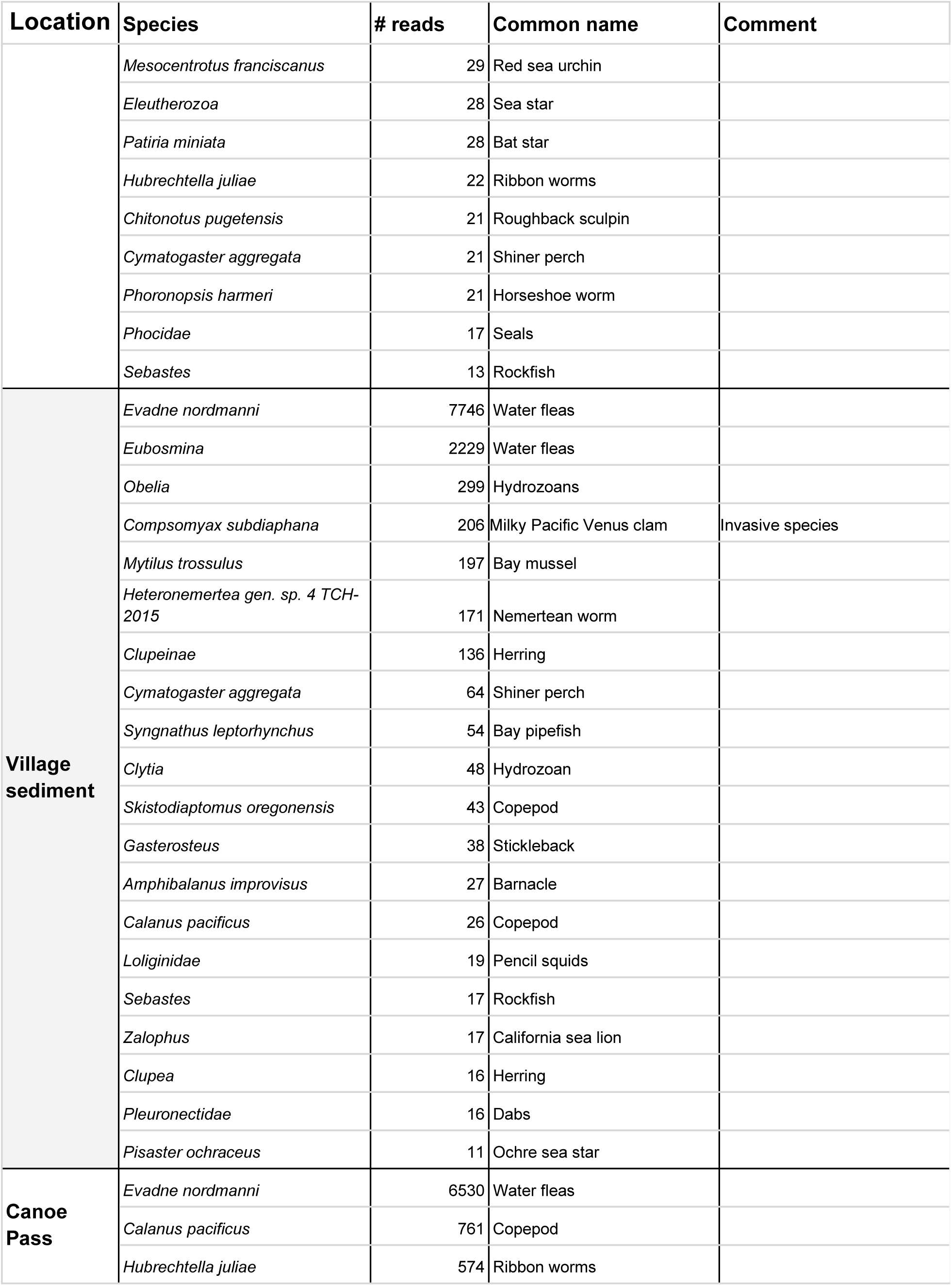
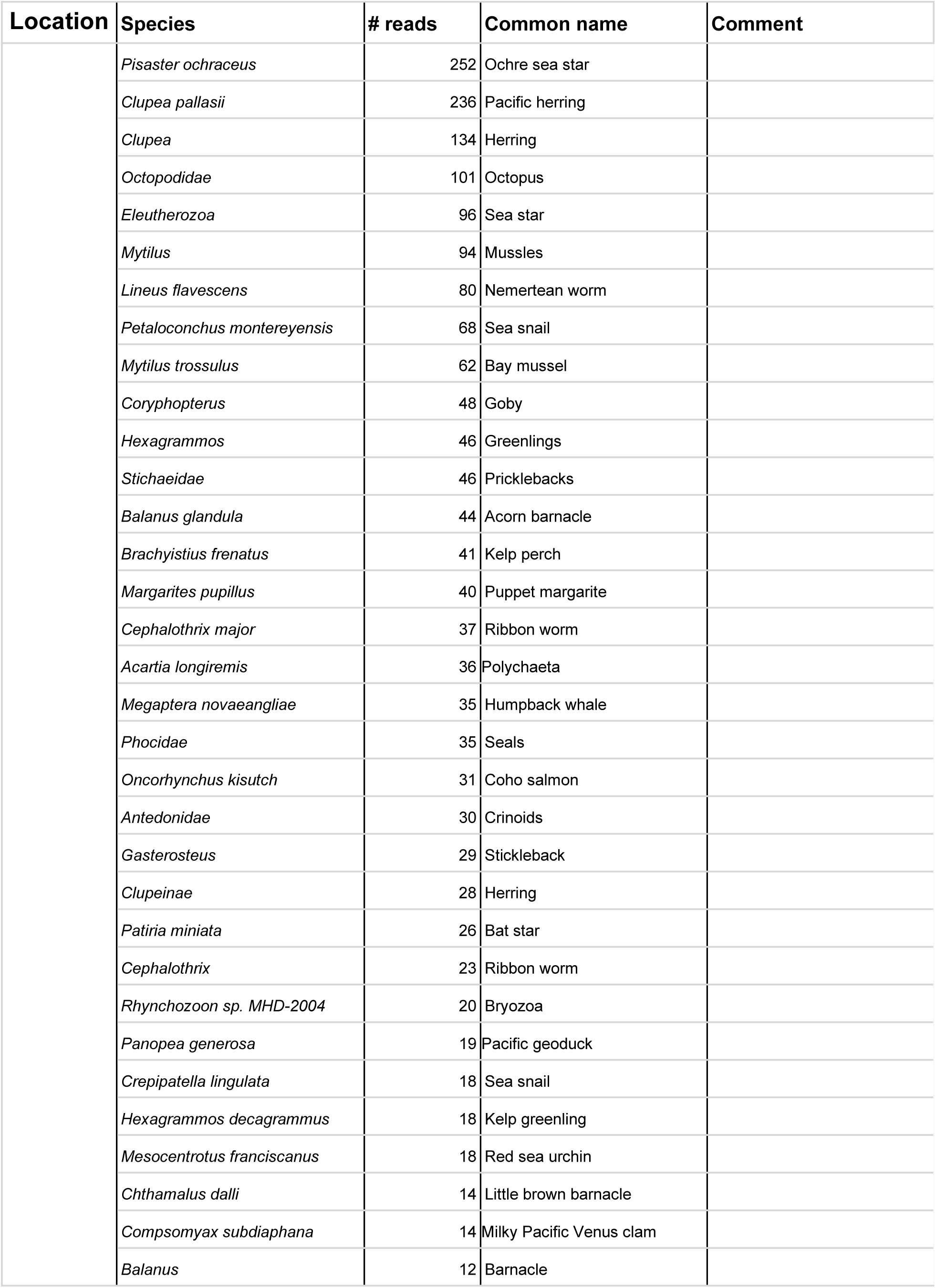
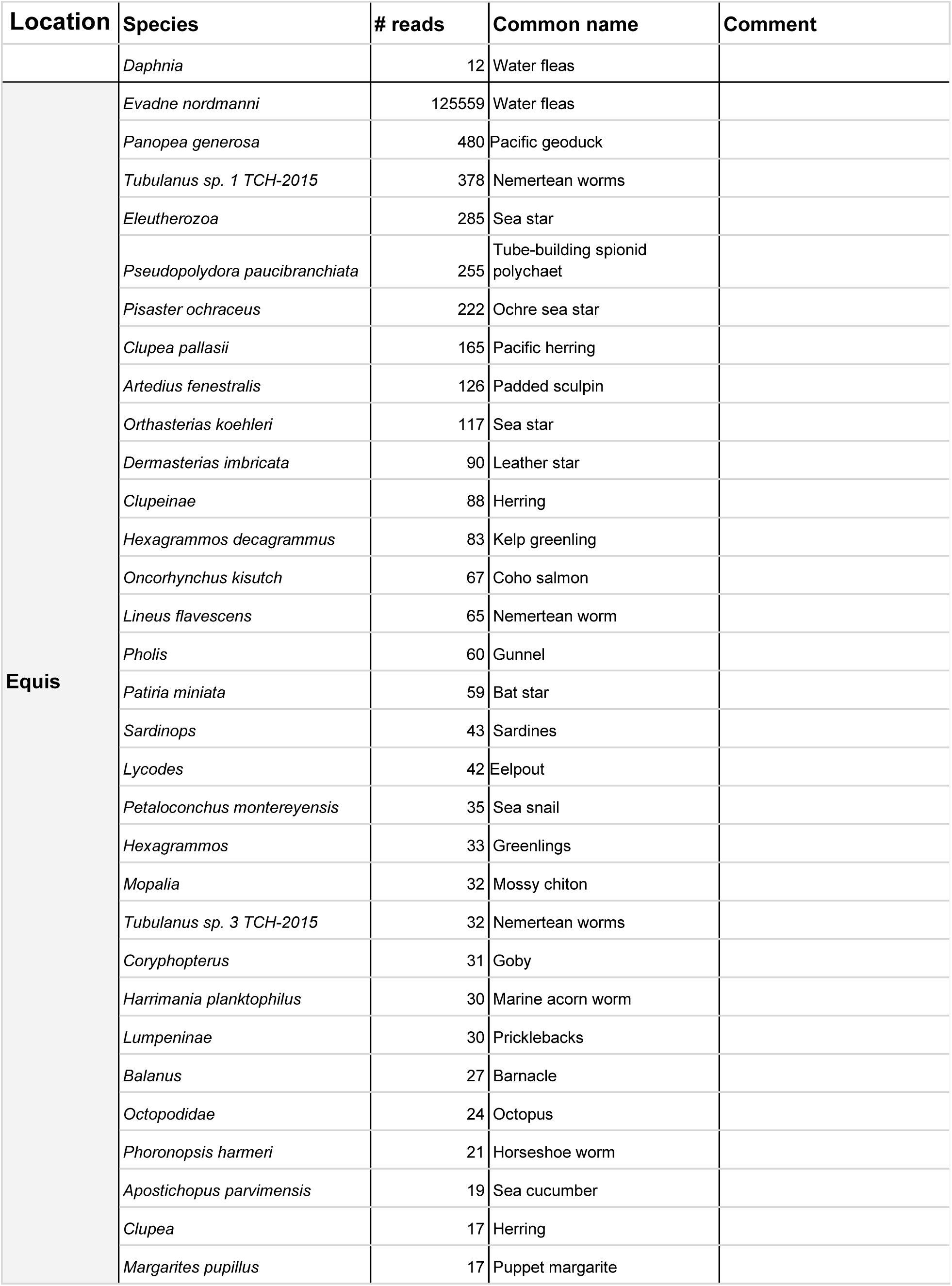
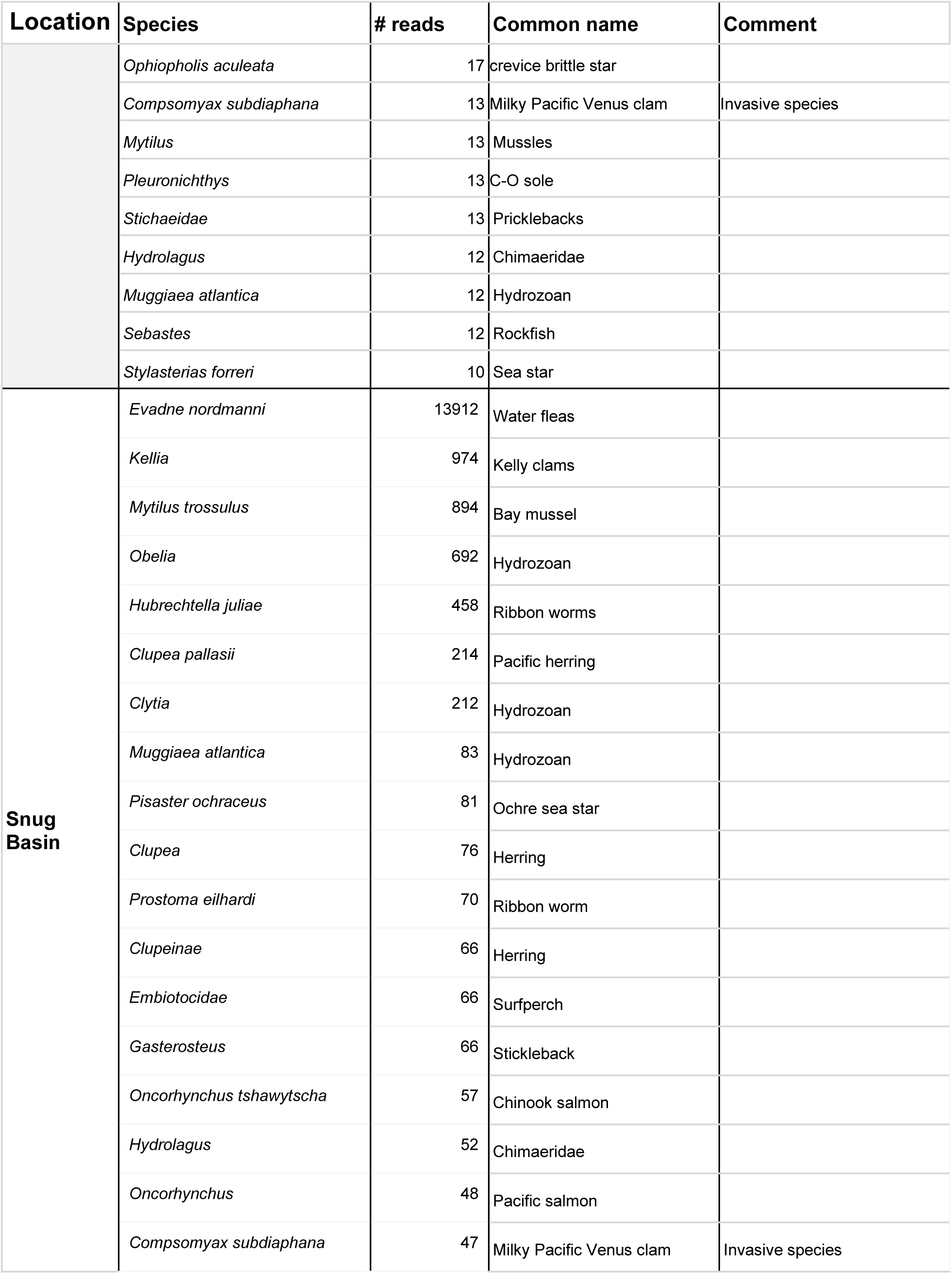
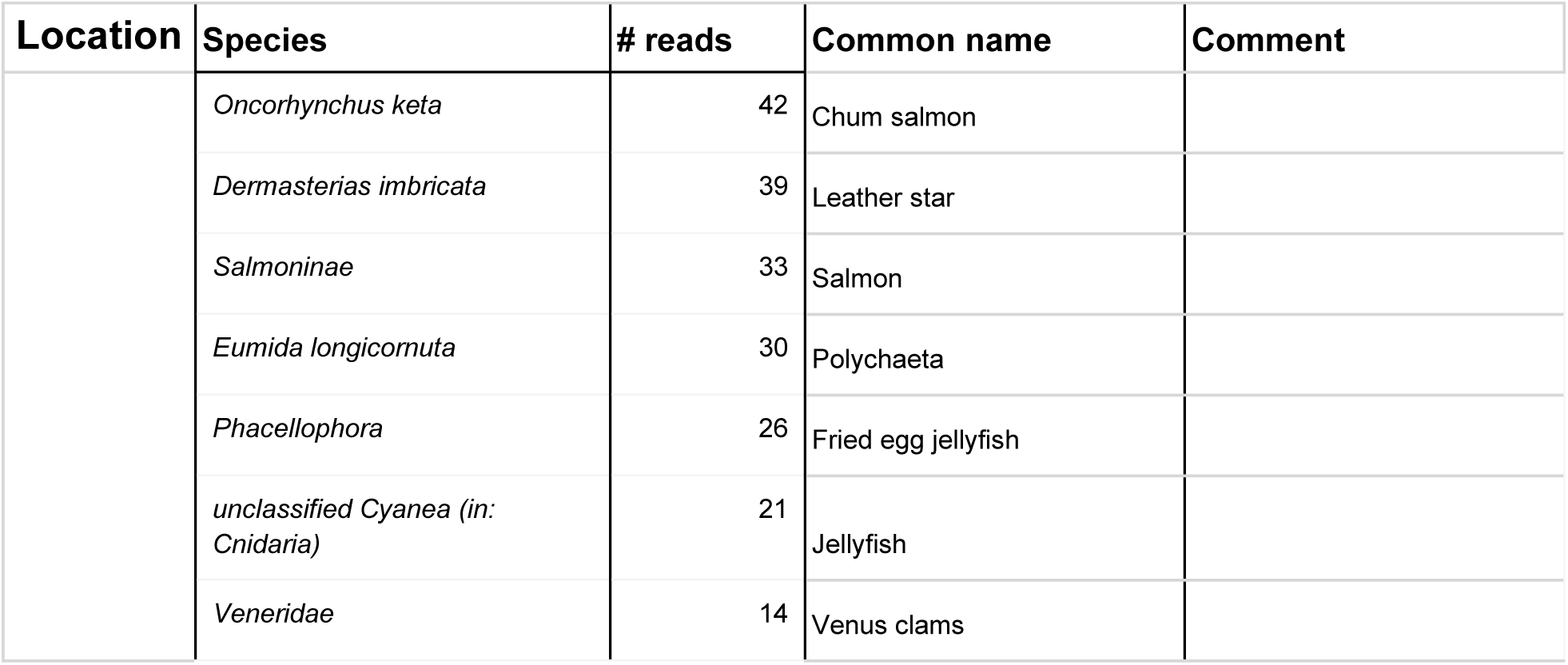
Detections in marine samples.

**Supplementary Figure 1:**
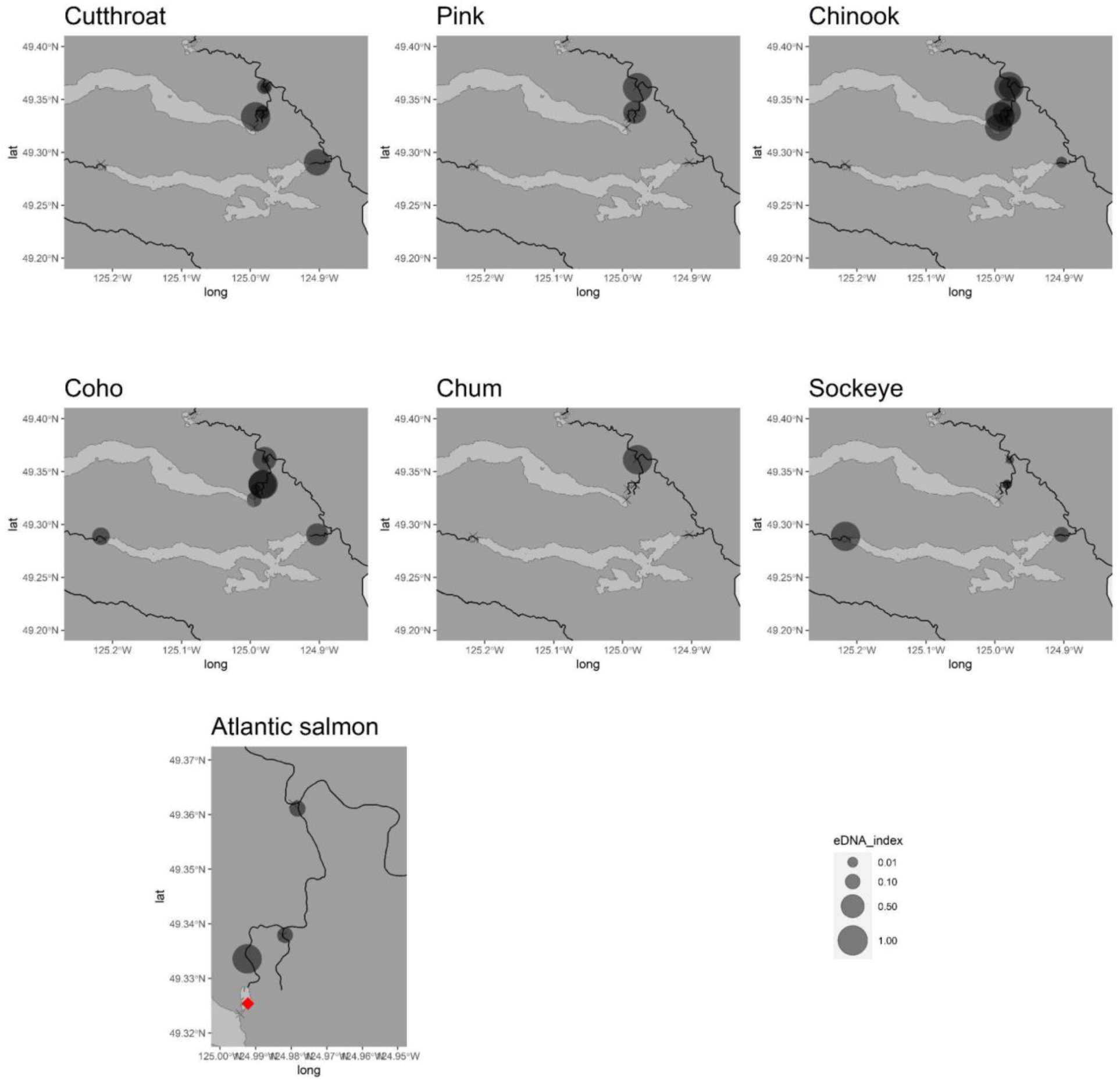
Salmonid detections in the Somass/Stamp watershed based on metabarcoding.

**Supplementary Figure 2:**
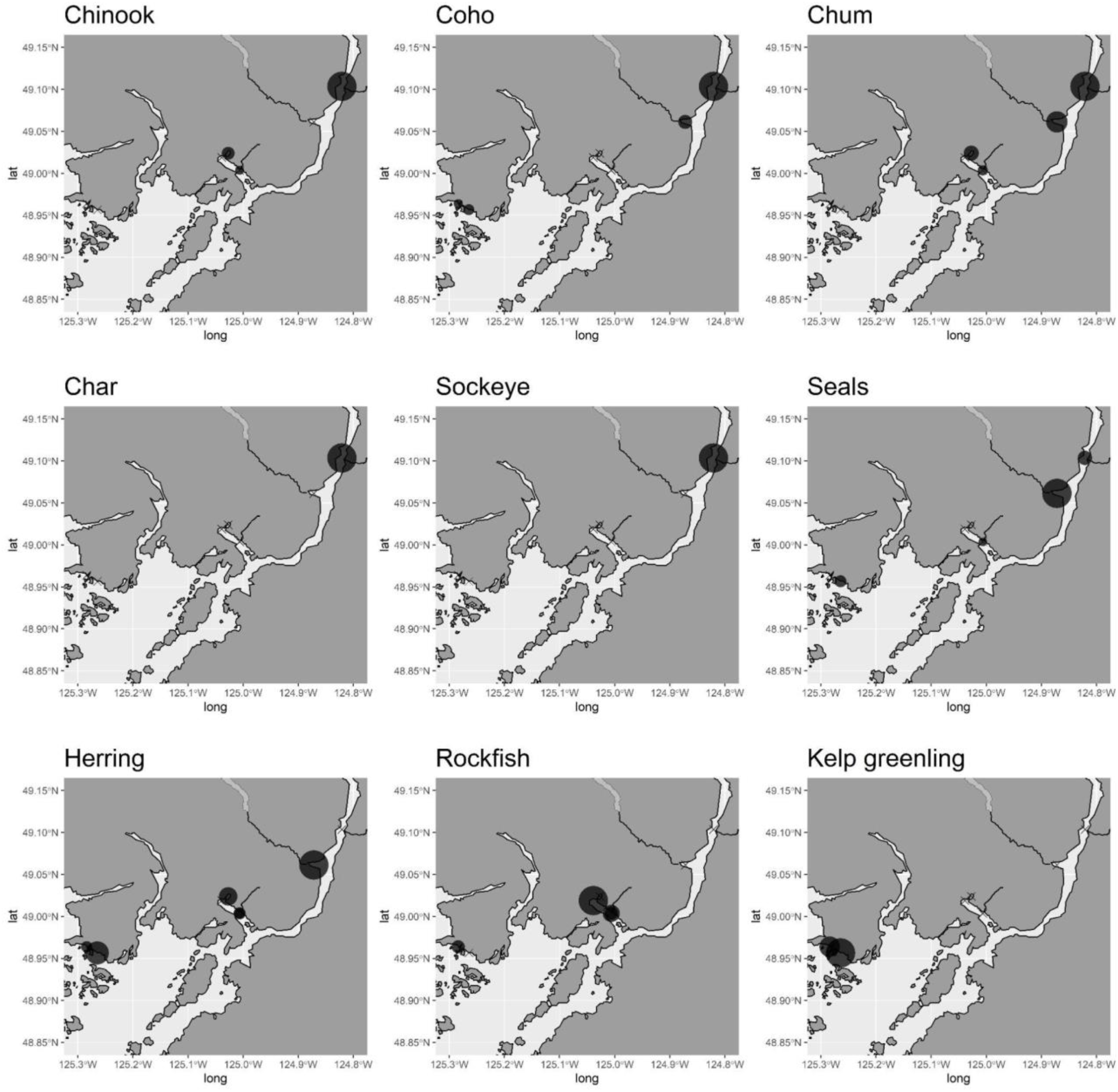
Vertebrate detections in the Alberni inlet and Broken islands based on metabarcoding.

**Supplementary Figure 3:**
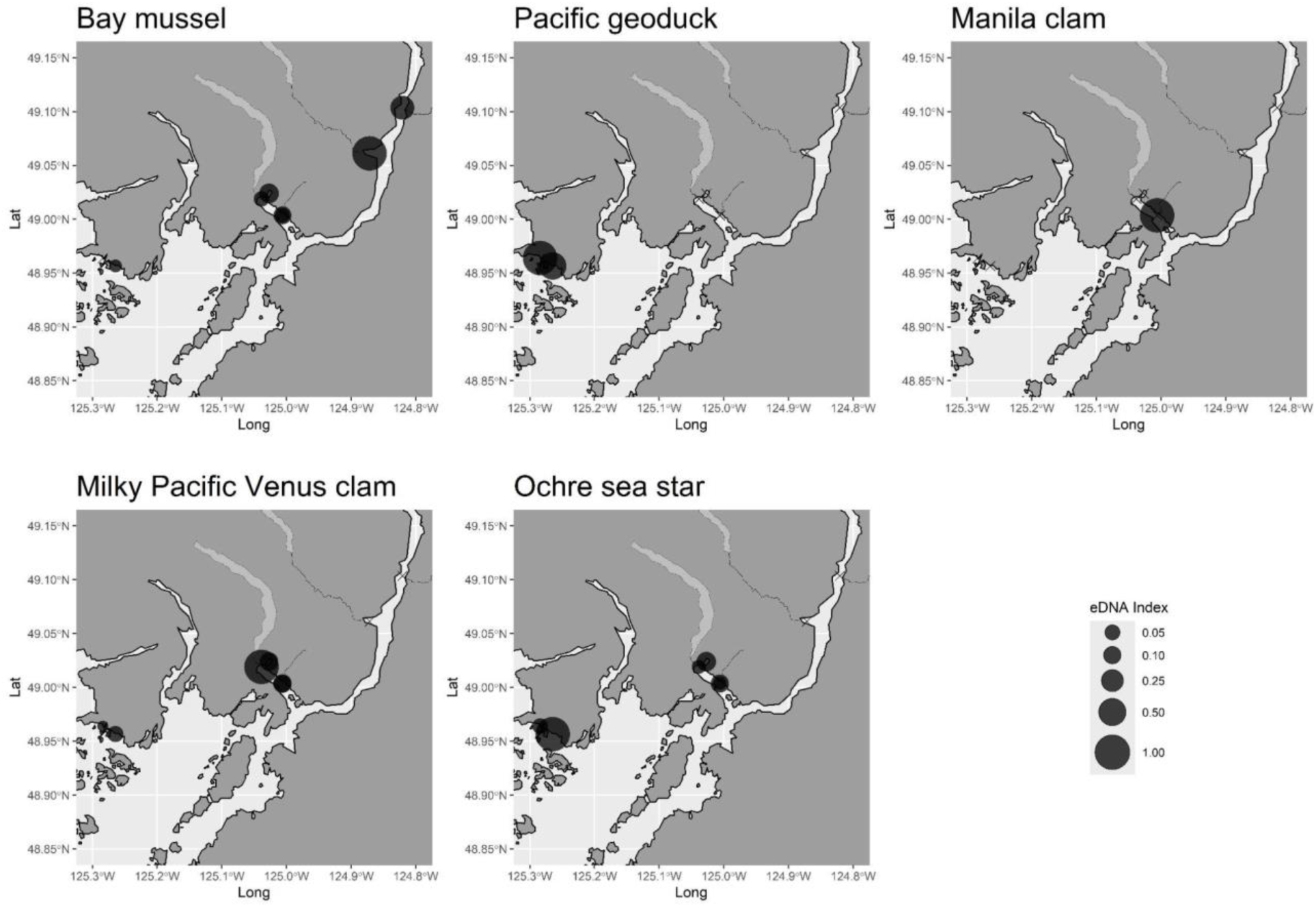
Invertebrate detections in the Alberni inlet and Broken islands based on metabarcoding.

